# Elevated Lactate in the AML Bone Marrow Microenvironment Polarizes Leukemia-Associated Macrophages via GPR81 Signaling

**DOI:** 10.1101/2023.11.13.566874

**Authors:** Celia A. Soto, Maggie L. Lesch, Jennifer L. Becker, Azmeer Sharipol, Amal Khan, Xenia L. Schafer, Michael W. Becker, Joshua C. Munger, Benjamin J. Frisch

## Abstract

Interactions between acute myeloid leukemia (AML) and the bone marrow microenvironment (BMME) are critical to leukemia progression and chemoresistance. In the solid tumor microenvironment, altered metabolite levels contribute to cancer progression. We performed a metabolomic analysis of AML patient bone marrow serum, revealing increased metabolites compared to age- and sex-matched controls. The most highly elevated metabolite in the AML BMME was lactate. Lactate signaling in solid tumors induces immunosuppressive tumor-associated macrophages and correlates with poor prognosis. This has not yet been studied in the leukemic BMME. Herein, we describe the role of lactate in the polarization of leukemia-associated macrophages (LAMs). Using a murine AML model of blast crisis chronic myelogenous leukemia (bcCML), we characterize the suppressive phenotype of LAMs by surface markers, transcriptomics, and cytokine profiling. Then, mice genetically lacking GPR81, the extracellular lactate receptor, were used to demonstrate GPR81 signaling as a mechanism of both the polarization of LAMs and the direct support of leukemia cells. Furthermore, elevated lactate diminished the function of hematopoietic progenitors and reduced stromal support for normal hematopoiesis. We report microenvironmental lactate as a mechanism of AML-induced immunosuppression and leukemic progression, thus identifying GPR81 signaling as an exciting and novel therapeutic target for treating this devastating disease.

## Key Points

- GPR81, the lactate receptor, is a mechanism of leukemia-associated macrophage (LAM) polarization to a suppressive phenotype.
- Targeting GPR81 reduces macrophage polarization, and has therapeutic potential for AML.

## Introduction

Acute myeloid leukemia (AML) is the most common acute leukemia in adults. It has a nearly 90% mortality rate at five years past diagnosis in the most affected group of greater than 65 years of age ^1^. AML is a hematologic malignancy initiated by genetic mutations in immature myeloid progenitor cells. During disease progression, leukemic cells proliferate in an uncontrolled manner leading to accumulation in the bone marrow (BM) and other tissues ^2^. Leukemic-initiated dysfunction of the bone marrow microenvironment (BMME) leads to a loss of normal hematopoiesis. This paucity of functional blood cells increases susceptibility to infection, hemorrhage, and BM failure, all critical factors in morbidity and mortality associated with AML ^3,4^. Chemotherapies initially reduce leukemic burden; however, relapse occurs in most patients due to surviving leukemia stem cells (LSCs) ^5–7^. With approximately 20,000 new cases annually in the United States [SEER, NIH] and an increasing global incidence ^8^, there is a clear, unmet need for novel treatments.

Signaling within the BMME directs hematopoietic stem cells (HSCs) to produce hematopoietic stem and progenitor cells (HSPCs). Multiple “HSC niche” cells regulate HSCs, including mesenchymal stromal cells (MSCs) ^9,10^, osteoblasts (OBs) ^11,12^, endothelial cells ^13–15^, and macrophages ^16^. HSC niche cells signal by contact-mediated mechanisms as well as secreted factors ^9,17–21^. The dysfunctional BMME not only loses support for normal hematopoiesis but also can gain support for leukemogenesis and leukemic progression ^22–24^. Furthermore, LSCs have been found to gain resistance to therapy by residing within the altered niche ^25,26^. Therefore, a thorough understanding of the microenvironment is needed to improve treatment.

Altered metabolite concentrations in the solid tumor microenvironment (TME) impede immune cells while supporting cancer ^27,28^. Thus far, the extracellular metabolites in the AML BMME have not been well-defined. A hallmark of cancer cells is energy production by aerobic glycolysis, called the “Warburg Effect”, even with functional mitochondrial oxidative phosphorylation (OXPHOS) ^29^. In a final step of glycolysis, lactate dehydrogenase (LDH) converts pyruvate to lactate. Extended glycolysis therefore demands secretion of excess lactate by cancer cells. Lactate concentrations have been reported to be elevated 5-30-fold in the TME and correlate with poor prognosis ^30,31^. Though heterogeneous within an individual, AML cells are known to upregulate both glycolysis and OXPHOS and are dependent on glycolysis for survival ^32,33^, and therefore would be expected to secrete lactate. Recently, increased lactate in AML BM has been reported ^34^, but is not yet well-documented. We hypothesized that metabolites accumulate in the AML BMME, including lactate, driving immunosuppression and leukemic progression.

Macrophages are critical to the stimulation of T cells during an immune response. Chronic signals in the TME, such as elevated lactate, alternatively polarize tumor-associated macrophages (TAMs); TAMs instead block T cells from targeting cancer cells. The presence of TAMs correlates with poor prognosis in multiple cancer types ^35–37^. An immune-suppressed BM is well-known in AML, yet lactate has never been directly connected to this. Leukemia-associated macrophages (LAMs) in AML have been reported to be alternatively activated, support leukemic transformation, and correlate with poor prognosis ^38,39^. Repolarization of macrophages toward a more pro-inflammatory phenotype impacts survival time in murine models of AML ^39,40^. Yet, little is known about the molecular mechanisms of how LAMs are polarized to this phenotype in AML. This research aimed to determine if elevated lactate in the BMME during myeloid leukemias contributes to suppressive macrophages.

## Methods

### Human bone marrow serum collection

Deidentified BM aspirates were collected from patients and immediately centrifuged to remove cells. The supernatant was quickly taken to storage at −80°C until use. Patients were eligible if they were diagnosed *de novo* for AML.

### Metabolomics by liquid chromatography mass spectrometry (LC-MS/MS)

LC-MS/MS was performed using an LC-20AD HPLC system (Shimadzu, Kyoto, Japan). Mass-spectrometric analyses were performed on a TSQ Quantum Ultra triple-quadrupole mass spectrometer running in multiple reaction monitoring mode (ThermoFisher Scientific). Peak heights for metabolite chromatograms were analyzed by using the Xcalibur software (RRID:SCR_014593). See **supplemental methods** for further details.

### Lactate measurements

Lactate concentrations were measured using a colorimetric lactate assay kit (Sigma MAK064). Absorbance was read on a BioTek Synergy 2 multi-mode microplate reader according to the manufacturer’s instructions.

### Flow cytometry and fluorescence-activated cell sorting (FACS)

Flow cytometric analyses were performed at the University of Rochester Wilmot Cancer Center on an LSRFortessa Cell Analyzer (BD Biosciences). FACS was performed at the Flow Cytometry Core at the University of Rochester Medical Center on a FACSAria II system (BD Biosciences) with an 85-micron nozzle at 4°C, using FACSDiva software (BD Biosciences, RRID:SCR_001456), then analyzed using FlowJo v10 software (BD Biosciences, RRID:SCR_008520). More details are listed in **supplemental**.

### RNA sequencing (RNAseq)

RNA sequencing and analysis were performed by the University of Rochester Genomics Research Center. See **supplemental methods** for further details.

### Cytokine profiling

A stromal monolayer of murine whole BM was grown to confluency in “complete” Minimum Essential Medium α (MEM α) (with nucleosides and L-glutamine, without ascorbic acid, Gibco A10490-01) with 10% heat-treated fetal bovine serum (htFBS) (Gibco 26140079) and 1x antibiotic-antimycotic (anti-anti) (Gibco 15240062)). Macrophages were sorted via FACS from the BM of bcCML or healthy mice, then cocultured with the stromal monolayer for 4 days to establish a microenvironment. Media was collected and stored immediately at −20°C until use. Proteome Profiler Mouse XL Cytokine Array (Bio-Techne (Minneapolis, MN, USA) R and D Systems ARY028) was used according to manufacturer protocol, then analyzed using the ChemiDoc MP Imaging System (Bio-Rad (Hercules, CA, USA)) and Image Lab software (Bio-Rad). Images were equally adjusted for background and the blot was qualitatively assessed for cytokine presence.

### Bone marrow-derived macrophage (BMDM) production

Whole BM was plated in a vented tissue culture (TC)-treated 75cm^2^ flask (NEST 708001) overnight in complete Dulbecco’s Modified Eagle Medium (DMEM) (Corning 10-013-CV). The next day, non-adherent cells (containing monocytes) were transferred to 12-well TC-treated culture dishes at 400,000 cells/well in 1 mL media with 25 ng/mL recombinant murine macrophage colony-stimulating factor (M-CSF) (PeproTech 315-02). Media was changed on day 4, when monocytes have adhered, and differentiation to BMDMs was complete by day 7.

### Macrophage polarization experiments

Cells were cultured in serum-fere media (without FBS) for 6 hours before treatment. Treatments were done in serum-free media for 48 hours, or the time indicated. Treatments included 10 mmol/L sodium L-lactate (Sigma), 100 ng/mL lipopolysaccharides (LPS) from E. coli O55:B5 (Sigma-Aldrich (St. Louis, MO, USA) L6529), and/or 5 ng/mL of recombinant murine interleukin (IL) −4 and −13 (IL-4, IL-13) (ILs) (ThermoFisher (Watham, MA, USA) PeproTech (Cranbury, NJ, USA) 214-14, 210-13). To remove cells for analysis, 0.25% trypsin-EDTA (ThermoFisher Gibco 25200056) was added for 5 min then cell scrapers were used.

### Colony forming unit (CFU-C) assays

Non-adherent cells were collected in a 15 mL conical tube, and 0.25% trypsin-EDTA was used on the adherent cells for five minutes, flushed with media, then added. Cells were pelleted by centrifugation for five minutes at 900 x g, then resuspended in 0.5 mL of MEM α. From here, both 1:20 and 1:100 dilutions were made in 2.5 mL of media, in separate Eppendorf tubes. Then, 0.2 mL of these dilutions were resuspended each in 2.5 mL aliquots of MethoCult (StemCell Technologies (Vancouver, CA) M3434) methylcellulose-containing media and immediately plated in duplicates of 1.2 mL into 35mm x 10mm sterile suspension culture dishes (Corning (Corning, NY, USA) 430588). Dishes were placed inside a sterile 150 mm x 25 mm dish (NEST (Woodbridge, NJ, USA) 715001), with an open dish of sterile water in the center to prevent drying. Then, were incubated for 14 days (5% CO_2_; 37C) and colonies were counted. Specific CFU experiments are described in **Supplemental Methods**.

## Results

### Extracellular metabolite levels are altered in the bone marrow of AML patients, including elevated lactate

To profile the metabolite levels of the AML BMME, we performed metabolomics on serum from BM biopsies of AML patients at diagnosis, as well as healthy age- and sex-matched controls (**Figure 1A**). The disease samples included different ages, sex, and mutational subtypes (**Supplemental Table S1**). AML BM displayed a general increase in extracellular metabolites (**Figures 1B** and **1C**). Six metabolites, listed in **Table 1**, were significantly altered during disease. Of these, lactate was the most highly elevated. Lactate concentrations were approximately 2-5-fold higher in AML BM, measured as 2-5 mmol/L (**Figure 1D**). *In vivo* concentrations are likely greater because BM biopsies are unavoidably diluted by peripheral blood. Also, we postulate that lactate concentrations are higher near AML cell-dense pockets, as the BM is spatially heterogeneous for concentrations of similar molecules and pH ^41^.

**Figure 1).**
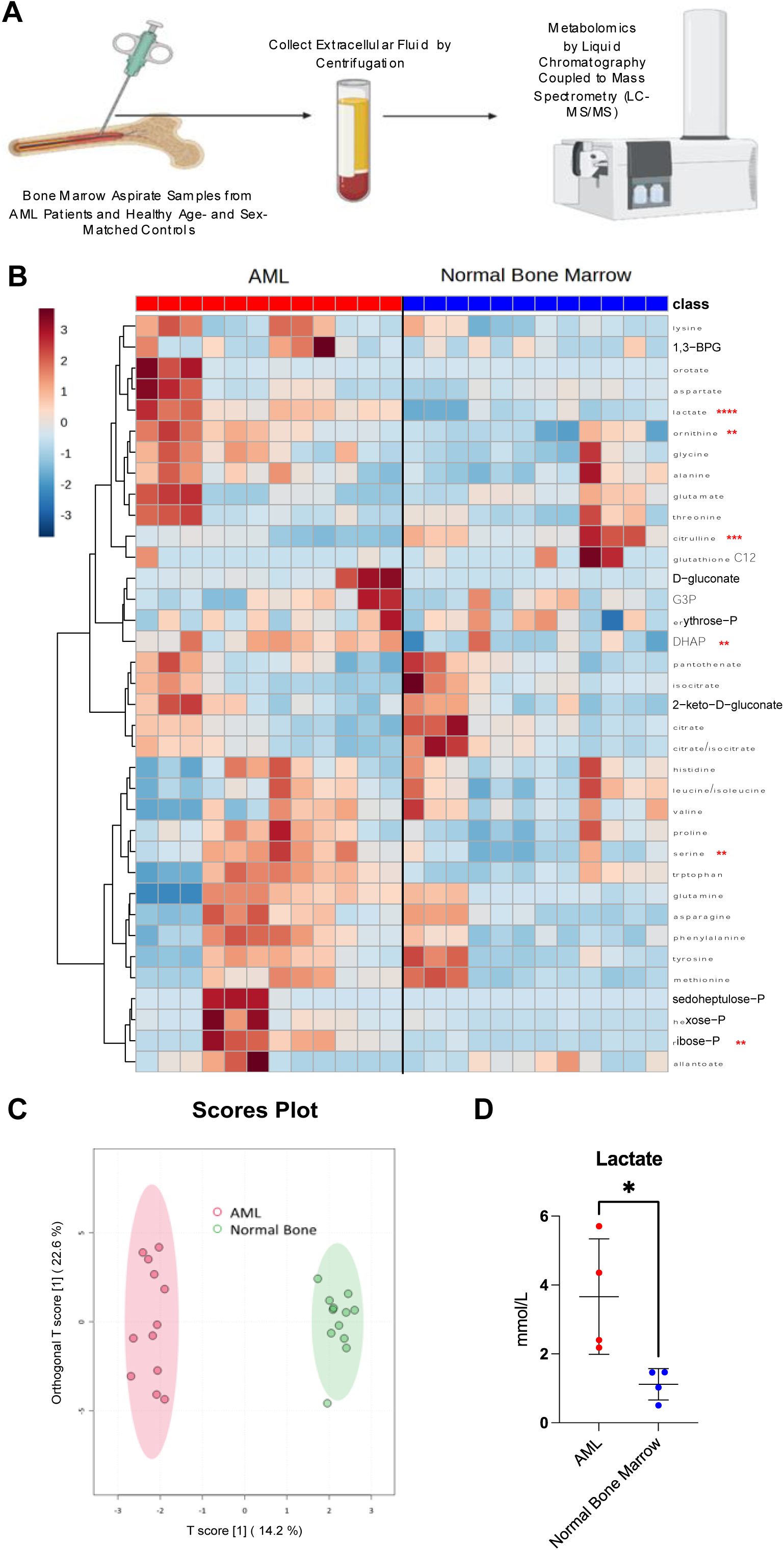
Lactate is elevated in the bone marrow microenvironment (BMME) during acute myeloid leukemia (AML). **A-C,** Metabolomics of bone marrow serum from AML patients or normal controls: Graphical depiction of procedure (**A**), heatmap showing the relative abundance of detectable metabolites (**B**), and scores plot of principal component analysis (**C**) (n = 4, in triplicates). **D,** Lactate concentration in AML and normal bone marrow (BM) serum (n = 4). Significance level determined by unpaired t test for **B** and **D** are indicated as: *, *P* < 0.05; **, *P* < 0.01; ***, *P* < 0.001; ****, *P* < 0.0001. Error bar indicates mean ± standard deviation (SD).

**Table 1:**
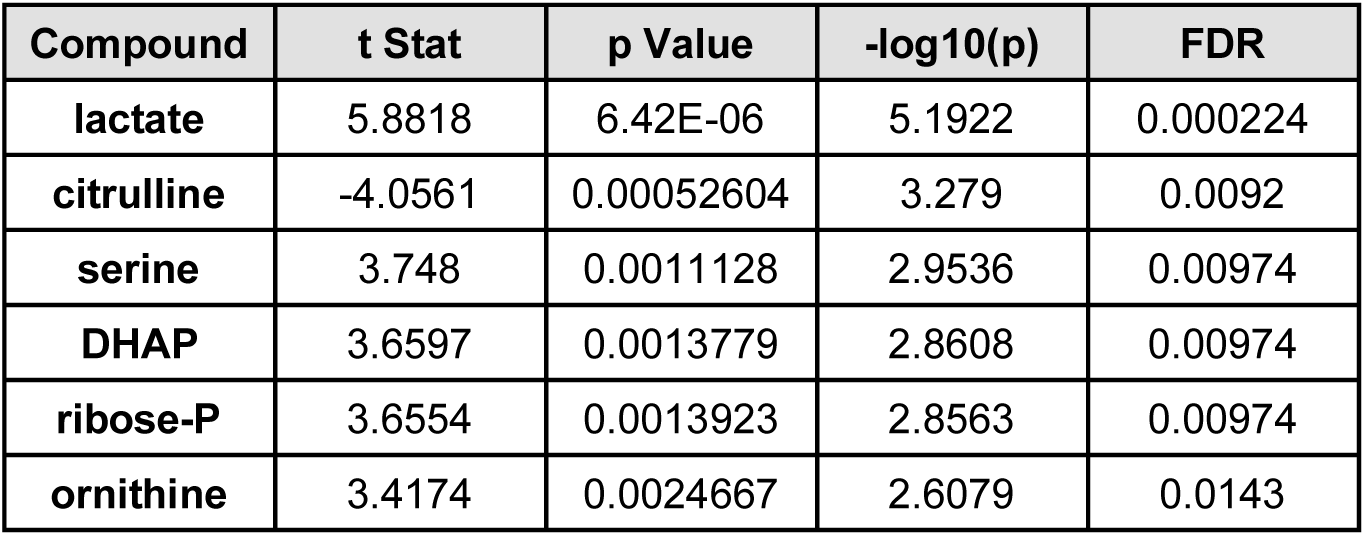
Significantly Altered Extracellular Metabolites in Human AML Bone Marrow.

Next, we aimed to study lactate signaling *in vivo*. We performed metabolomics on BM extracellular fluid from a previously described murine AML model, blast-crisis chronic myelogenous leukemia (bcCML) (**Supplemental Figure 1A**) ^23,42^. BM metabolites and lactate were elevated in bcCML BM (**Supplemental Figure 1B** and **Supplemental Table S2**). The lactate increase was relative to AML (**Supplemental Figure 1C**), demonstrating that this model is well-suited.

### Leukemia-associated macrophages (LAMs) are alternatively activated to a suppressive phenotype

We hypothesized that lactate is immunosuppressive in AML, through macrophage polarization. We profiled the activation phenotype of BM macrophages (Ly-6C^-^, Ly-6G^-^, CD45^+^, F4/80^+^) from leukemic mice compared to nonleukemic (NL) controls (**Figure 2A**). Leukemic cells were distinguished from nonleukemic myeloid cells by co-expression of green fluorescent protein (GFP). Well-described macrophage activation markers were assayed: CD38 and major histocompatibility complex class II (MHCII) as classic/proinflammatory markers ^43,44^, or early growth response protein 2 (EGR2) and macrophage mannose receptor (MR/CD206) as alternative/suppressive markers ^45,46^. The SPICE 6.1 program (NIH) was used to quantify the frequency of phenotype subsets. This unbiased approach identified a CD206^+^ subset of LAMs that are enriched in disease (**Figure 2B**), a marker of suppressive TAMs associated with poor prognosis ^47–49^.

**Figure 2).**
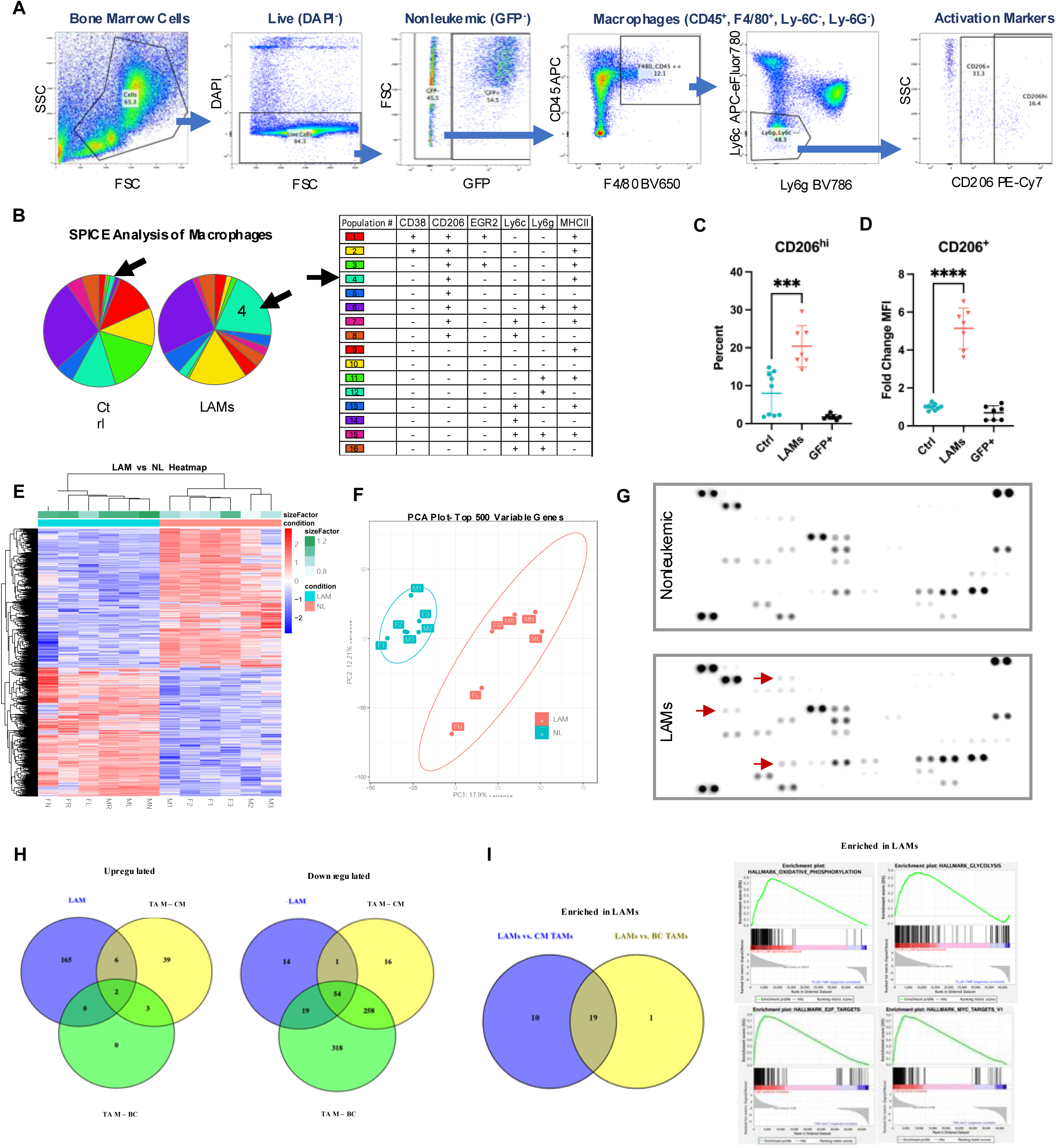
Leukemia-associated macrophage (LAMs) display an alternatively activated, suppressive phenotype. **A,** Gating scheme for flow cytometric analysis of polarization markers on murine macrophages. **B,** SPICE analysis showing frequency of macrophage subpopulations from healthy controls or late-stage bcCML (n = 5), arrows indicate population enriched in disease. **C** and **D,** Nonleukemic control macrophages (Ctrl), leukemia-associated macrophages (LAMs), or leukemia-derived (GFP^+^) macrophages (n = 7) from bcCML BM. Frequency of CD206^hi^ (**C**) and expression level by mean fluorescence intensity (MFI) of CD206 on CD206^+^ (**D**). **E** and **F,** Bulk RNA sequencing of LAMs vs. macrophages from nonleukemic (NL) controls. Heatmap of differentially expressed genes (**E**) and principal component analysis (PCA) of the top 500 variable genes (**F**). **G,** Macrophages were sorted from nonleukemic or leukemic mice and cultured for four days (n = 2), then media was profiled for cytokines. Representative example, arrows indicate a qualitative difference. **H,** Venn diagrams displaying the number of GO pathways significantly upregulated or downregulated by murine cancer-associated macrophages compared to each study’s own healthy controls: LAMs (n = 6) or tumor-associated macrophages (TAM) from colorectal liver metastasis (CM) (n = 5) or breast cancer (BC) (n = 3). **I,** Hallmark gene sets enriched in LAMs compared directly to TAMs, determined by gene set enrichment analysis (GSEA): Venn diagram displaying numbers of significantly enriched gene sets, and representative enrichment plots of top significant sets. Significance levels determined by one-way ANOVA for **C** and **D,** are indicated as: ***, *P* < 0.001; ****, *P* < 0.0001. Error bar indicates mean ± SD. Significance for **I** was determined by GSEA as an FDR q-value of < 0.25.

By the time of late-stage disease (≥50% leukemic cells in BM), there was an increase in the frequency of CD206^hi^ LAMs (**Figure 2C**) and expression level of CD206 (**Figure 2D**). This was only observed in the non-leukemia-derived (GFP^-^) cells displaying macrophage markers. The frequency of MHCII^+^ macrophages was lower in bcCML, while CD38 and EGR2 were unchanged (**Supplemental Figures 2A-C**). High CD206 and low MHCII on LAMs indicates a shift towards a suppressive phenotype. Increased CD206 was detectable by early-stage disease (7-10% leukemic cells in the BM) (**Supplemental Figures 2D** and **2E**).

We further examined the phenotype of LAMs by transcriptomics. The transcriptome of LAMs was distinct from NL macrophages (**Figures 2E** and **2F**). The top upregulated Gene Ontology (GO) pathways indicate altered interactions with immune and hematopoietic cells (**Supplemental Figure 2F**). Top downregulated pathways were related to neutrophils and cell cycle control (**Supplemental Figure 2G**). Then, by proteome profiling *in vitro*, we detected several cytokines exclusively expressed by LAMs (**Figure 2G** and **Table 2**). These are implicated in solid tumors for an immunosuppressive microenvironment (CCL12, CCL6, PCSK9) ^50,51^, tumor cell proliferation and invasion (CXCL10) ^52^, immune cell chemotaxis (CCL12/MCP-5, CXCL10) ^50,52^, and tumor growth and metastasis (CXCL5/LIX, MMP3, proprotein convertase 9 (PCSK9)) ^53–55^. These data confirm that LAMs are polarized to a suppressive phenotype.

**Table 2:**
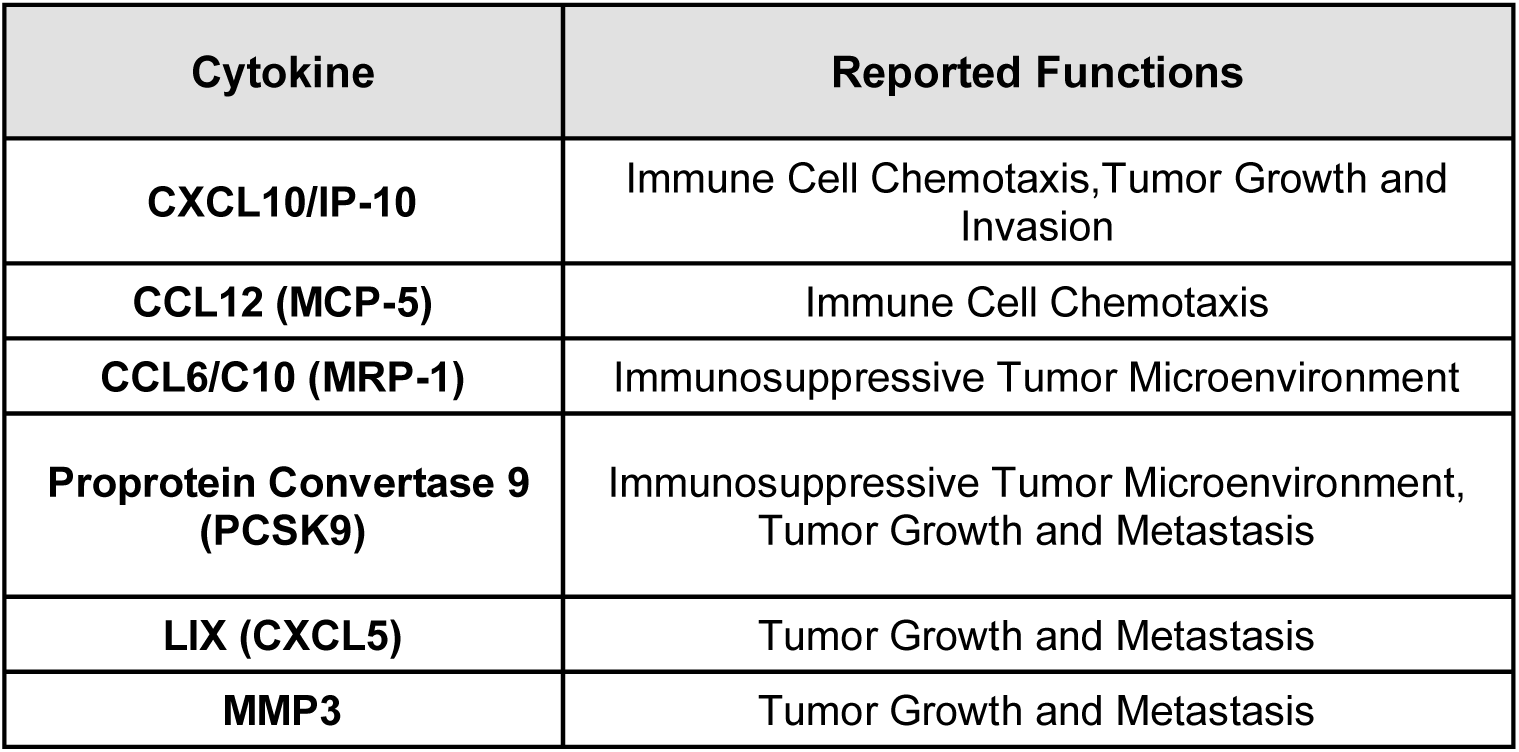
Cytokines Upregulated in LAM Cocultures and Reported Functions in Cancer.

To our knowledge, this is the first published murine transcriptomic dataset of macrophages from leukemic BM. As such, we asked how similar LAMs are to TAMs from other types of cancers. LAMs were compared to murine F4/80^+^ TAMs from solid tumor models of colorectal metastasis (CM) ^56^ and breast cancer (BC) ^57^ (NIH National Center for Biotechnology Information (NCBI) Gene Expression Omnibus (GEO) accession viewer IDs GSE206211 and GSE126268). Pathway analysis identified GO elements upregulated/downregulated in both LAMs and TAMs compared to their internal control macrophages from noncancerous tissue (**Figure 2H**). Common upregulated elements are listed in **Table 3**, and downregulated elements in **Supplemental Table S3**.

**Table 3:**
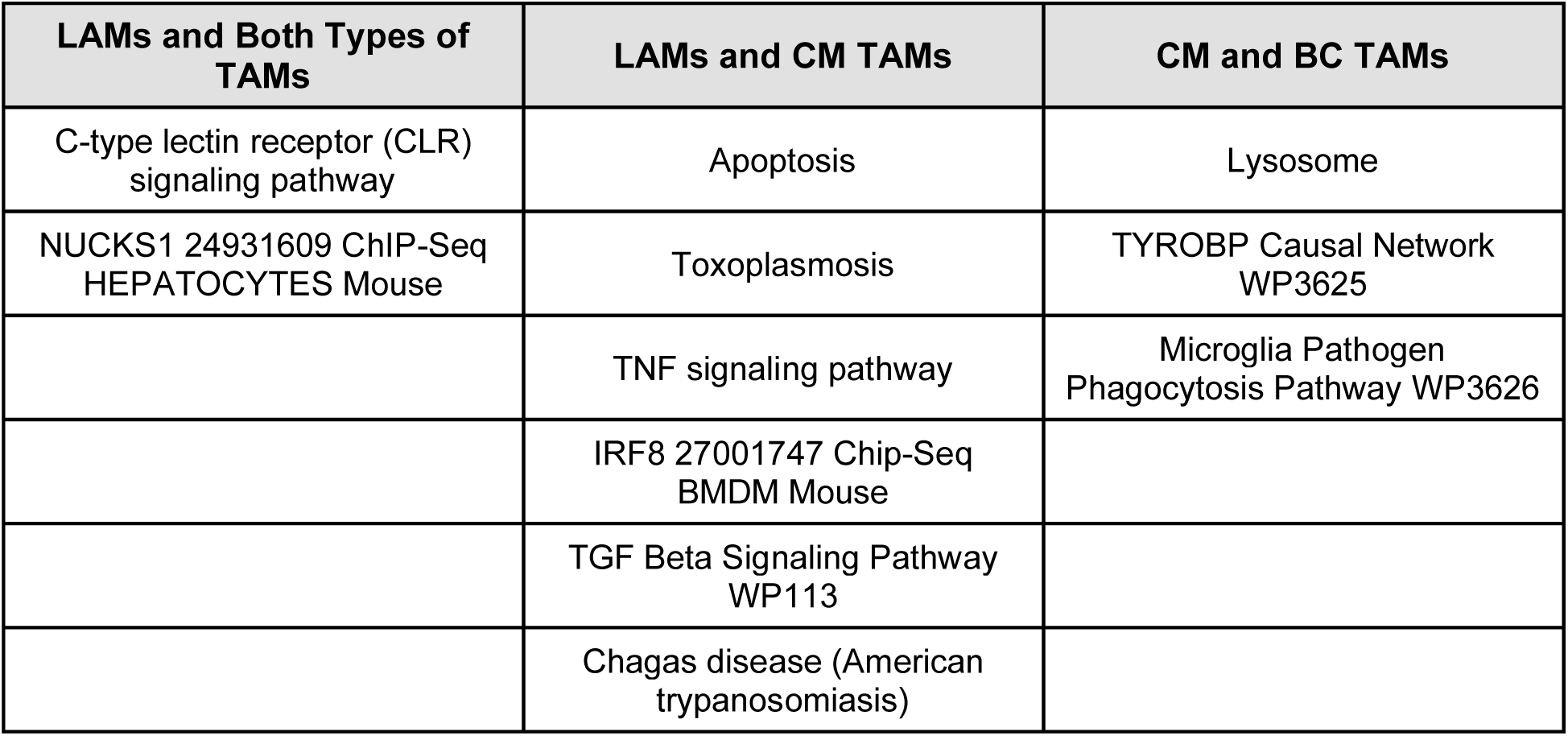
Common Upregulated Elements in LAMs and TAMs.

Two elements were common to LAMs and both types of TAMs. One was C-type lectin receptor (CLR) signaling, which functions in immunomodulation by recognizing polysaccharides from pathogens ^58^. A notable CLR is CD206, which was elevated on bcCML LAMs. The other common upregulated element of LAMs and TAMs was nuclear casein kinase and cyclin-dependent kinase substrate 1 (NUCKS1), a nuclear DNA binding protein that is ubiquitously expressed in mice and humans. NUCKS1 functions in signal transduction of the cell cycle and DNA damage response, and regulates inflammation through NF-κβ mediated cytokine expression ^59^. Though overexpression in various cancers has been reported ^60,61^, NUCKS1 in macrophages has not yet been studied. Additionally, LAMs and CM TAMs upregulated tumor necrosis factor (TNF) and transforming growth factor β (TGF-β) signaling, two key pathways related to T cell immunomodulation and the permissive cancer microenvironment ^62–64^, and interferon regulatory factor 8 (IRF8), involved in chronic inflammation, myeloid differentiation, and the activation of macrophages ^65^. There were a greater number of common downregulated elements, listed in **Supplemental Table S3**, including p53 signaling which is important for the resolution of alternative phenotype ^66^, and others largely involved with cell cycle, DNA replication, and mitosis.

Then, we directly compared LAMs and TAMs by performing gene set enrichment analyses (GSEA). “Hallmark” gene sets from the Human Molecular Signatures Database (MSigDB) represent pathways with homology in humans. LAMs were enriched for metabolic, cell cycle, and stress related pathways compared to both TAM types (**Figure 2I**), listed in **Supplemental Table S4**. Though the pathway analysis show that overall cell cycle is downregulated in LAMs compared to healthy macrophages, this may be less so than in their solid tumor counterparts. Instead, both TAM types were enriched for angiogenesis epithelial to mesenchymal transition pathways more relevant to solid tumors (**Supplemental Figure 2H**), and BC TAMs for several inflammatory signaling pathways listed in **Supplemental Table S5**. In all, LAMs display a CD206^hi^, immune suppressive phenotype like TAMs, yet with key differences that highlight them as a unique population in need of further study.

### GPR81 signaling is a mechanism of lactate-induced LAM polarization

To isolate the influence of lactate on polarization, we utilized bone marrow-derived macrophages (BMDMs) treated *in vitro* with physiologically relevant levels of lactate. As positive controls, we applied lipopolysaccharide (LPS) as proinflammatory/classic stimuli, or interleukins (IL) IL-4 and IL-13 as alternative stimuli ^67–70^. IL-4/13 are well-described Type 2 helper T (Th2) cytokines that persist in the AML BMME ^71^. Polarization to CD206^hi^ BMDMs increased synergistically when lactate was combined with IL-4/13 (**Figure 3A** and **3B**).

**Figure 3).**
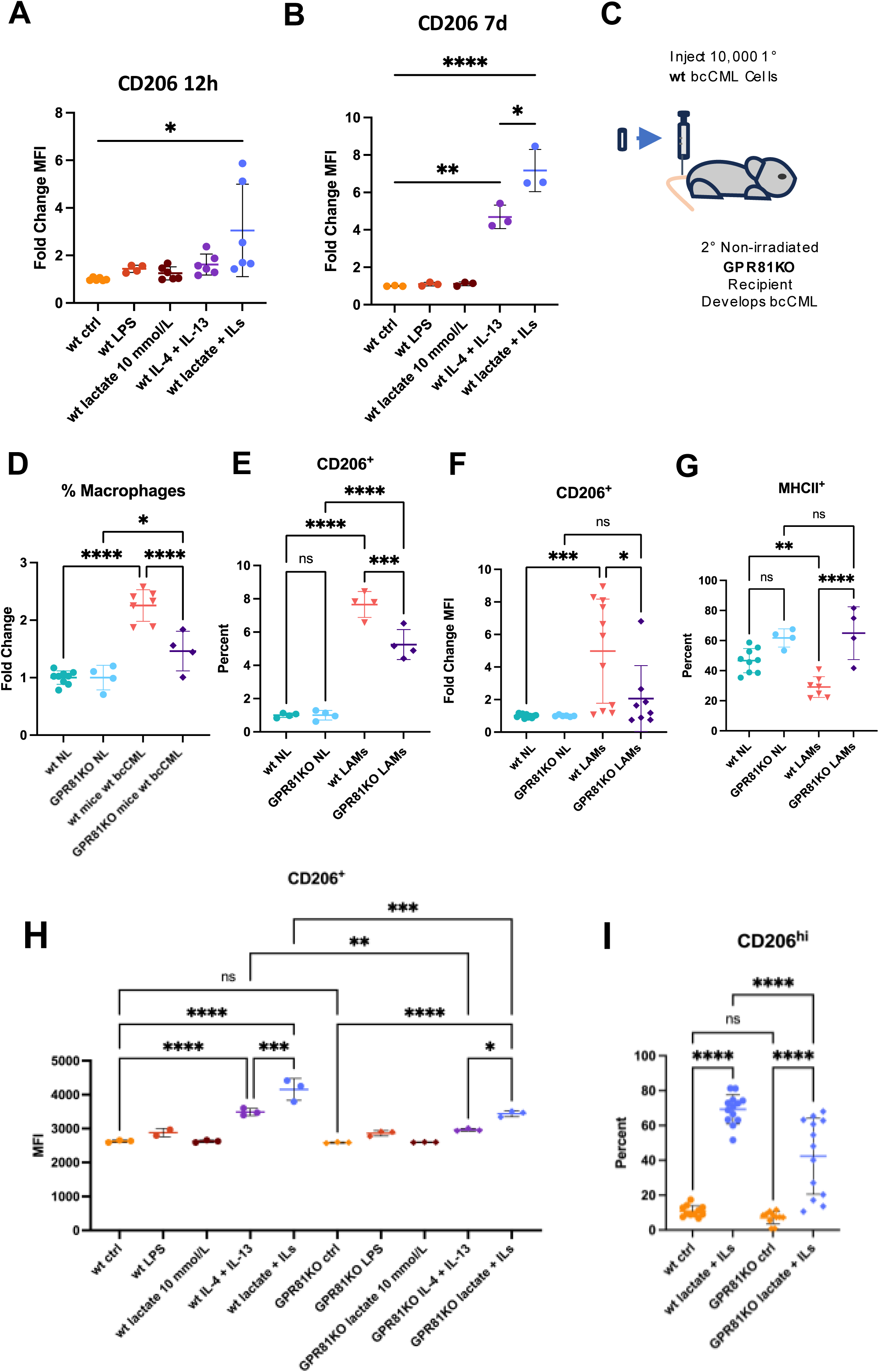
Lactate-GPR81 signaling contributes to leukemia-associated macrophage (LAM) polarization. **A** and **B,** Fold change expression level of CD206 on bone marrow-derived macrophages (BMDMs) *in vitro* treated with lipopolysaccharide (LPS), lactate, and/or IL-4/-13 (ILs), for for 12 hours (**A**) (n = 6) or 7 days (**B**) (n = 3). **C,** Depiction of the initiation of bcCML in a GPR81KO BMME. **D-G,** Flow cytometric analysis of nonleukemic (GFP^-^) bone marrow (BM) cells from bcCML in wt or GPR81KO mice, and nonleukemic (NL) controls. Frequency of macrophages in the BM relative to NL controls of the same genetic background (**D**) (n = 4-7). BM macrophages: frequency of CD206^+^ (**E**), expression level of CD206 relative to NL controls of the same genetic background (**F**), frequency of MHCII^+^ (**G**) (n = 4-11). **H,** Expression of CD206 on wt or GPR81KO BMDMs treated with LPS, ILs, and/or lactate (n = 3). **I,** Frequency of CD206^hi^ wt or GPR81KO BMDMs with or without polarization by ILs and lactate. Significance levels determined by one-way ANOVA for **A-B, D-G,** and **H-I** are indicated as: ns, not significant; *, *P* < 0.05; **, *P* < 0.01; ***, *P* < 0.001; ****, *P* < 0.0001. Error bar indicates mean ± SD.

Generally, the activation of macrophages to a classic/proinflammatory phenotype is marked by upregulated expression of inducible nitric oxide synthase (iNOS/Nos2), which mediates the cytotoxic production of NO to assist pathogen killing and phagocytosis. Instead, alternative/suppressive macrophages express Arginase 1 (Arg1), involved in depleting the substrate of iNOS, L-arginase ^72,73^, to resolve an immune response. CD206^hi^ BMDMs expressed *Arg1*, and not *iNOS* (**Supplemental Figures 3A** and **3B**). These findings support that CD206^hi^ LAMs are functionally suppressive.

Next, we aimed to determine the specific lactate signaling pathway affecting LAMs. Lactate is an extracellular ligand to G-protein-coupled hydroxycarboxylic acid receptor 1 (GPR81/HCAR1), the cell-surface “lactate sensor” ^74^. GPR81 has been linked to the pathophysiology of cancer and the immune-suppressed TME. For example, GPR81 signaling inhibits antigen-presenting cells in lung and breast cancers, proinflammatory NF-κB signaling in macrophages, and inflammasome activity ^75–77^. Also, cross-membrane lactate transport is coordinated through monocarboxylate transporters (MCT)- 1 and −4 ^78,79^. Intracellularly, lactate activates pathways such as hypoxia-inducible factor-1 (HIF-1), which has been connected to TAM polarization ^35^, and can be consumed as metabolic fuel via the citric acid cycle ^80^. While inhibiting MCT1/4 or LDH has been studied as a therapeutic approach for targeting lactate in AML cells ^32,81^, the hematopoietic system also relies on these key proteins for homeostasis, leading to off-target effects. However, GPR81 has not yet been studied in AML.

To investigate lactate-GPR81 signaling as a mechanism of polarization, we utilized mice genetically lacking GPR81 (GPR81KO). Wt bcCML was initiated in GPR81KO mice (**Figure 3C**). The increased frequency of BM macrophages during bcCML was partially reversed in the GPR81KO mice (**Figure 3D**). Fewer LAMs in the GPR81KO BMME expressed CD206, and at lower amounts (**Figures 3E** and **3F**). This indicates that GPR81-lactate signaling contributes to LAM polarization. Also, the frequency of MHCII^+^ macrophages increased (**Figure 3G**). By late-stage disease, GPR81KO mice displayed a reduced leukemic burden in the peripheral blood and spleen (**Supplemental Figures 3C-E**). Next, we repeated BMDM polarization *in vitro*; GPR81KO BMDMs were not as polarized as wt BMDMs from the same stimuli (**Figures 3H** and **3I**), and *Arg1* expression was decreased relative to CD206 expression (**Supplemental Figure 3F**). Therefore, targeting GPR81 is a novel therapeutic option to reduce suppressive LAMs.

### Elevated lactate is harmful to the hematopoietic bone marrow microenvironment

Unfortunately, most current treatment options for AML exacerbate alterations to the BMME, and the subsequent loss of normal hematopoiesis that lead to fatal complications of the disease. Therefore, novel therapeutic targets for AML must also consider effects on the hematopoietic BMME. We considered that excess lactate in the BM may also be harmful to normal hematopoiesis. As we have previously reported, bcCML presents with an increased percentage HSPCs (LSK), as well as multipotent progenitors (MPP): megakaryocyte-biased MPP2, myeloid-biased MPP3, lymphoid-primed MPP4, as well as MSCs, a key stromal cell type for the maintenance of HSPCs (**Supplemental Figures 4A-G**) (see **Supplemental Methods Table SM2** for markers) ^23,82^. Still, mature blood cell populations are lost as leukemia progresses, indicating dysfunction in the progenitors. Therefore, we investigated whether high BM lactate or LAMs affect HSPC maintenance.

To determine if lactate reduces the hematopoietic potential of HSPCs, colony-forming (CFU-C) assays were performed ^83,84^. HSPCs lost colony-forming potential when treated with increasing lactate (**Figure 4A**). We then asked if coculture with HSC-niche supportive stromal cells could increase the maintenance of HSPCs in increased lactate, however, CFU-Cs were still reduced (**Supplemental Figures 4H and 4I**). Since we have previously reported that aged macrophages can also alter the colony-forming potential of HSPCs ^85^, we tested the addition of LAMs to the HSPC cocultures rather than lactate (**Supplemental Figure 4J**). HSPCs showed reduced CFU-Cs when cocultured with LAMs compared to healthy macrophages (**Figure 4B**), suggesting that LAMs provide altered hematopoietic maintenance signals.

**Figure 4).**
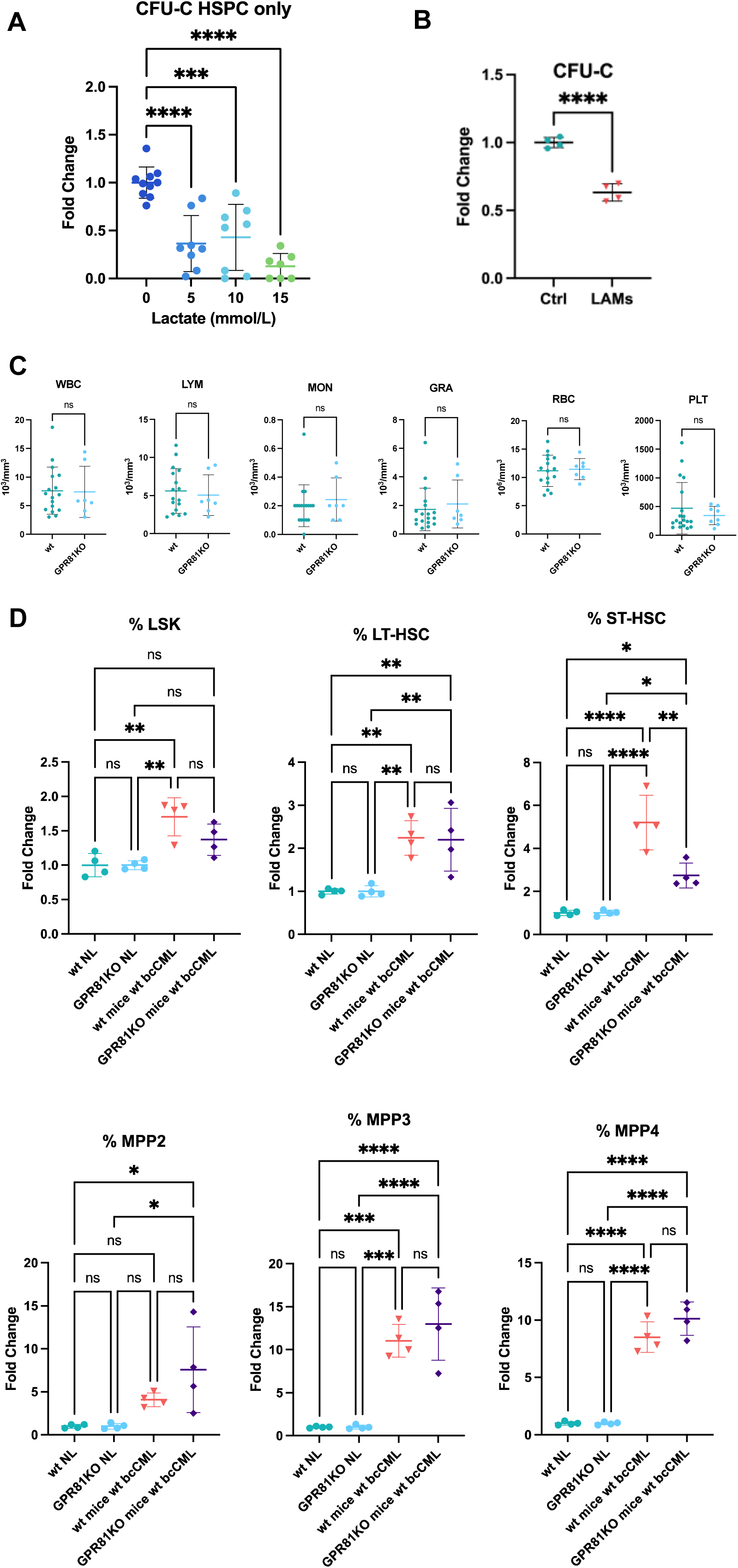
Excess lactate negatively effects the function of hematopoietic stem and progenitor cells (HSPC) and support, partially regulate d by GPR81. **A,** HSPCs treated with lactate for 72 hours, fold change colony forming unites (CFU-C) relative to the 0 mmol/L lactate control group (n = 7-10). **B,** HSPCs cocultured with LAMs or healthy control (Ctrl) macrophages for four days: fold change CFU-Cs relative to control group (n = 4). **D,** Hematopoietic progenitors’ frequency in the bone marrow, from bcCML in wt or GPR81KO mice, relative to the nonleukemic (NL) control of the same genetic background: hematopoietic stem and progenitor cells (HSPCs/LSK), long-term hematopoietic stem cell (HSC), short-term HSC, multipotent progenitor (MPP) subsets MPP2, MPP3, and MPP4 (n = 4). Significance levels determined by one-way ANOVA for **A** and D, or by unpaired t test for **B** are indicated as: ns, not significant; *, *P* < 0.05; **, *P* < 0.01; ***, *P* < 0.001; ****, *P* < 0.0001. Error bar indicates mean ± SD.

We posited that GPR81 may be largely dispensable to normal hematopoiesis because nonleukemic, adult GPR81KO mice have appropriate ratios of mature blood cells (**Figure 4C**). Next, we investigated whether hematopoietic progenitors are impacted by GPR81 signaling during pathologic conditions. An expansion of LSKs and ST-HSCs was abrogated when bcCML was initiated in GPR81KO mice, and other progenitor populations were unchanged (**Figure 4D**). Though, GPR81KO HSPCs still showed a moderate reduction of CFU-C loss when treated with lactate, compared to wild type (wt) (**Supplemental Figures 4K** and **4L**).

We also asked whether lactate alters key HSC niche stromal cells: MSCs and OBs. Lactate treatment reduced differentiation and self-renewal of stromal cultures (**Supplemental Figures 4M-O**). This is consistent with a loss of functional OBs and bone volume that we have previously reported in AML ^23^. These results highlight multiple damaging effects of elevated lactate on critical components of hematopoiesis in the BM. Altogether, these data show that reducing GPR81 signaling has a positive impact on hematopoietic progenitors, without harmful effects on normal hematopoiesis or stromal support.

### Lactate-GPR81 signaling drives leukemia cell growth and self-repopulation

Finally, we investigated whether GPR81 signaling impacts myeloid leukemia cells, as it is known to be crucial for the survival of other types of cancer cells ^86^. To specify, GPR81 enhances growth/survival pathways, immunoevasion by upregulating programmed death-ligand 1 (PD-L1), and chemoresistance via compound export (ABCB1 transporter) ^86–89^.

GPR81KO bcCML was initiated in GPR81KO mice (double knockout (DKO)) or in wt mice (**Supplemental Figure 5A**). Leukemic burden was largely reduced in the BM, peripheral blood, and spleen by the time point of late-stage disease compared to wt controls (**Figures 5A-D**). There was a delayed expansion of engrafted cells (**Supplemental Figures 5B-D**), and the time to progression to >50% leukemic cells in the BM was significantly increased in GPR81 DKO (**Figure 5E**).

**Figure 5).**
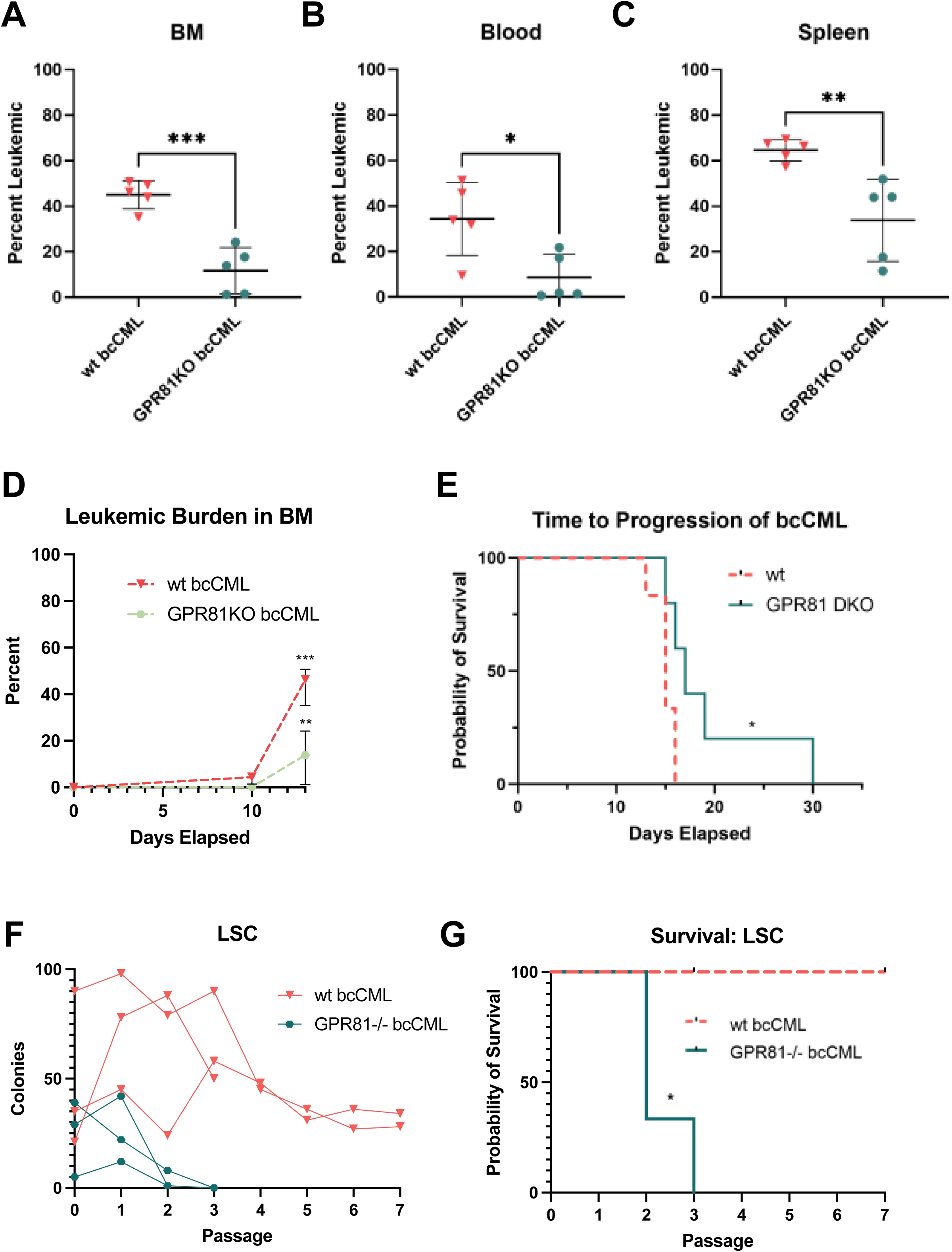
GPR81 signaling drives leukemic cell expansion rate and leukemia stem cell (LSC) self-renewal. **A-C,** Leukemic burden in the bone marrow (BM) (**B**), peripheral blood (**C**), and spleen (**D**), of wt or GPR81KO bcCML, at the timepoint of late-stage disease in wt (n = 5). **D,** Leukemic burden in the BM over time (n = 5-7). **E,** Time to progression to late-stage disease in wt or GPR81 DKO bcCML (n = 5). **F** and **G,** LSC repopulation, number of colonies at each passage (**F**), and survival curve (**G**) (n = 3, in duplicates). Significance levels determined by unpaired t tests for **A-D**, and log-rank (Mantel-Cox) test for **E** and **G**, are indicated as: *, *P* < 0.05; **, *P* < 0.01; ***, *P* < 0.001. Error bar indicates mean ± SD.

As residual LSCs after chemotherapy are the primary reason for relapse in patients ^5,25^, we experimentally determined the impact of GPR81 signaling on LSC self-repopulation. When serial passaging in methylcellulose-containing media, GPR81KO bcCML cells produced fewer colonies on average at P0 and P1 and lost repopulating capacity by P2-P3, while wt bcCML cells continued to repopulate colonies to at least P7 (**Figures 5F** and **5G**). These data display the importance of GPR81 signaling to the rapid growth and self-repopulation of leukemia cells. Altogether, we report that GPR81 is a mechanism of LAM polarization, and a potential therapeutic target for LAMs and leukemia cells that spares normal hematopoietic function.

## Discussion

Current AML chemotherapies commonly lead to relapse and damage the BM. Effective immunotherapies for myeloid leukemias are still under development, and pose the challenge of shared antigens with the myeloid immune cells. Herein, we report elevated levels of metabolites in the BMME during AML, defining elevated lactate as a critical driver of AML-induced macrophage polarization and leukemia progression as visually demonstrated in **Figure 6**.

**Figure 6).**
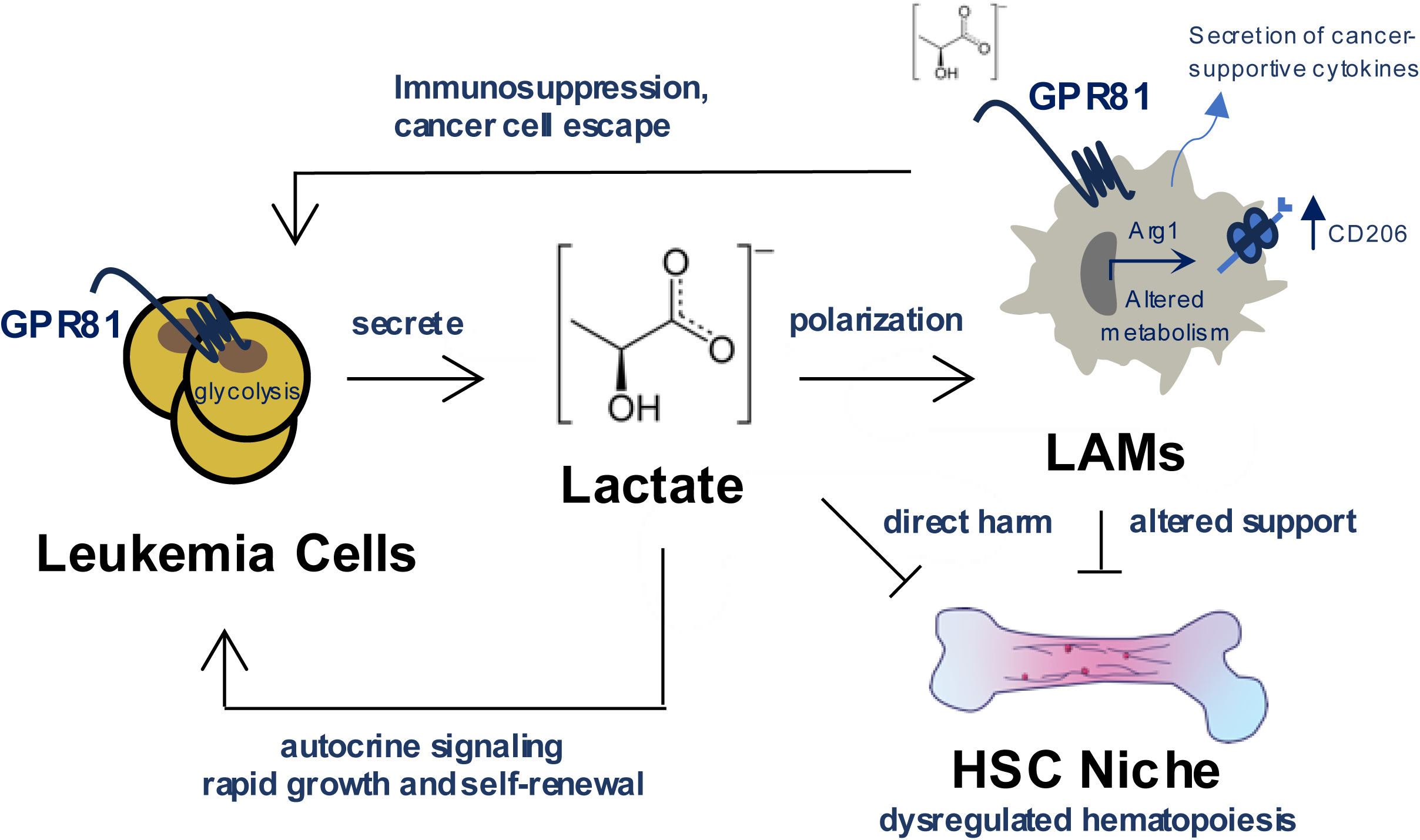
Effects of elevated lactate in the AML bone marrow microenvironment (BMME). Lactate is elevated in the AML BMME. This polarizes LAMs to a suppressive phenotype, characterized by increased CD206 expression, secretion of cancer-supportive cytokines, expression of Arg1, and altered metabolism. Increased lactate also harms normal hematopoiesis both directly and through altered support by LAMs. Autocrine lactate signaling supports the growth and repopulation of leukemia cells. The lactate receptor GPR81 is a mechanism of LAM polarization, and is implicated in these pathologic mechanisms of AML progression.

Our results highlight the potential of GPR81 as a novel therapeutic target for both the leukemic suppression of the immune system and for AML cell self-repopulation, with mild preservative effects on the hematopoietic BMME. Additionally, as lactate production is a hallmark of cancer, findings on the mechanisms of lactate signaling within the BMME are potentially applicable to multiple malignancies with BM involvement, including additional types of leukemia as well as bone metastases of solid tumors.

We identify lactate as the most elevated extracellular metabolite in the AML BMME. Lactate has been reported to have the strongest prognostic risk value of metabolites detected in the serum from cytogenetically normal AML patients ^90^. Our results highlight the lactate receptor GPR81 as a mechanism of LAM polarization. LAMs were CD206^hi^, displaying transcriptome and cytokine profiles associated with immunosuppression and cancer support. CD206 has been recently suggested as another prognostic factor for AML ^91^ and is induced on monocytes cocultured with AML blasts ^72^. The LAM-secreted cytokines we identified herein may be further investigated for their role in AML, particularly in the suppression of a cytotoxic T cell response. Increasing the pro-inflammatory capabilities of macrophages has the potential to increase the efficacy of chemotherapies as well as immunotherapies, such as CAR-T cell therapy.

A recent study by Weinhäuser et al. performed single-cell RNA sequencing on human AML-associated macrophages (AAMs). They reported that AAMs displayed alternative polarization, decreased phagocytosis, upregulated mitochondrial function, and overexpression of CD206 ^38^. The AAMs correlated with poor prognosis in a cohort of MDS patients and influenced engraftment of patient-derived xenografts, showing the clinical importance of suppressive macrophages. Further, the similarity of murine LAMs to AAMs highlights the translatability of our results to human AML ^38^.

Our research also supports the further study of T cell suppression by lactate in AML. High lactate can contribute to expression of PD-L1 on tumor cells, CD8+ T cell exhaustion, and suppression of MHCII^hi^ immune cells ^75,92^. Interestingly, T_Regs_ are more resistant to lactate-mediated inhibition than other T cell types ^93^, and are increased in AML patients ^94^. Furthermore, alternative macrophages induce chemotaxis/differentiation of T_Regs_ to the TME ^51,95,96^. At the hematopoietic niche, T_regs_ provide an immune-suppressed niche where LSCs may escape immune attack ^97^. The impact of lactate, GPR81 signaling, and LAMs on T cell subsets in AML has not yet been studied.

In non-leukemic mice, experimental knockout of GPR81 did not cause negative effects on hematopoiesis or stromal cell support. To the contrary, in leukemic mice loss of GPR81 exhibited a protective effect on the expansion of HSPC populations that may lead to their exhaustion. These novel and exciting findings include both murine and human data on lactate levels in the BM, highlighting the relevance to human disease. Further research will help elucidate the role of GPR81 in a human environment and solidify the potential of GPR81 as a therapeutic target.

Taken together, our findings demonstrate the lactate receptor GPR81 as a mechanism for the polarization of suppressive leukemia-associated macrophages and as a driver of AML cell growth. This research suggests that targeting GPR81 signaling in the BMME during AML may be a selective and well-tolerated therapeutic option to prevent LSC repopulation and rescue microenvironmental dysfunction.

## Acknowledgments

We kindly thank Dr. Vadivel Ganapathy, Ph.D. at Texas Tech University for gifting the GPR81KO mice. We would like to acknowledge the Genomics Research Center at the University of Rochester Medical Center for RNA sequencing and analysis, the flow cytometry core for FACS. Financial support for this project came from the Wilmot Cancer Institute Research Development Funding Program Pilot Award to Dr. Frisch and Dr. Munger, an American Cancer Society Grant RSG-22-165-01-MM to Dr. Frisch, and from a National Research Service Award (NRSA) Institutional Research Training Grant (T32) 5T32AR076950-03 from the National Institutes of Health (NIH) to Dr. Soto through the Rochester Musculoskeletal (ROCMSK) Training Program at the University of Rochester Center for Musculoskeletal Research (CMSR).

## Authors’ contributions

**CAS:** Conceptualization, experiments, data analysis, methodology, writing, and editing. **MLL, AS, and AK** assisted with experiments. **JLB** performed RNA sequencing analyses comparisons to online databases. **XLS** assisted with LC/MS-MS training and analysis. **MWB** collected human samples and contributed to conceptualization. **JCM:** Conceptualization, methodology, and data analysis. **BJF:** Conceptualization, data analysis, methodology, reviewing/editing, and oversight of the project. All authors read and approved the final manuscript.

