## Supplemental Figures for "Elevated Lactate in the AML Bone Marrow Microenvironment Polarizes Leukemia-Associated Macrophages via GPR81 Signaling"

#### Supplemental Fig. 1

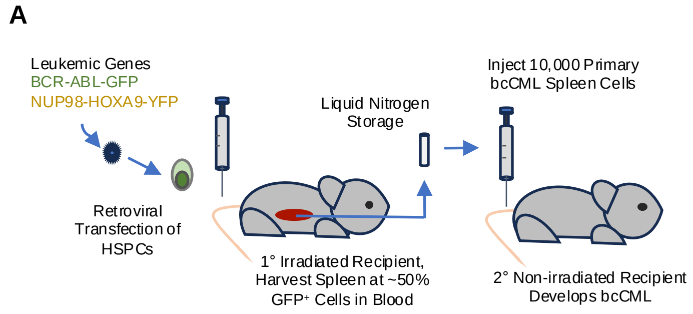


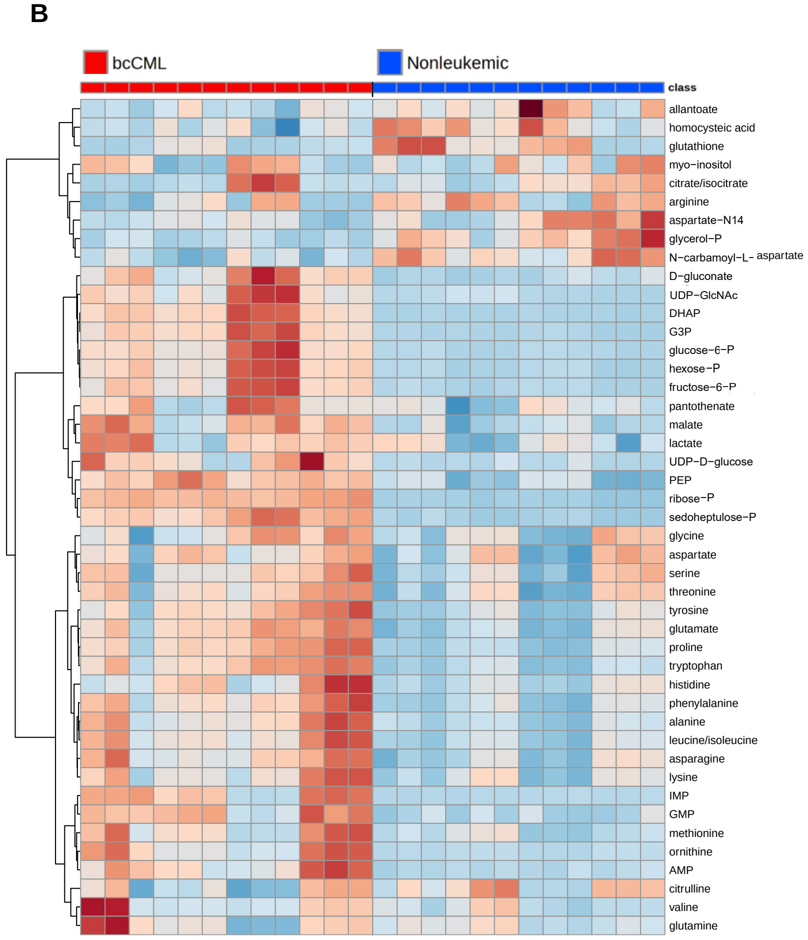


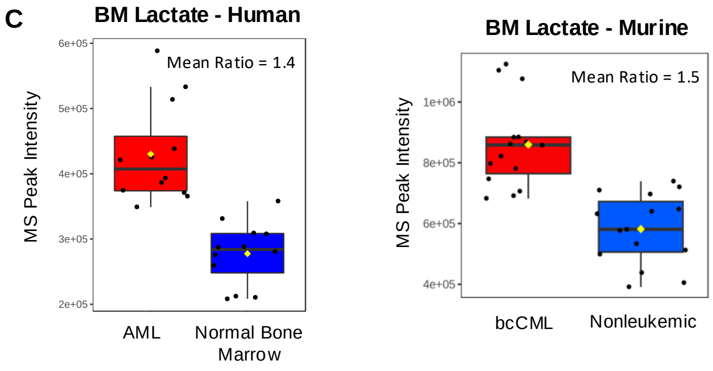

**Supplemental Fig. 2**

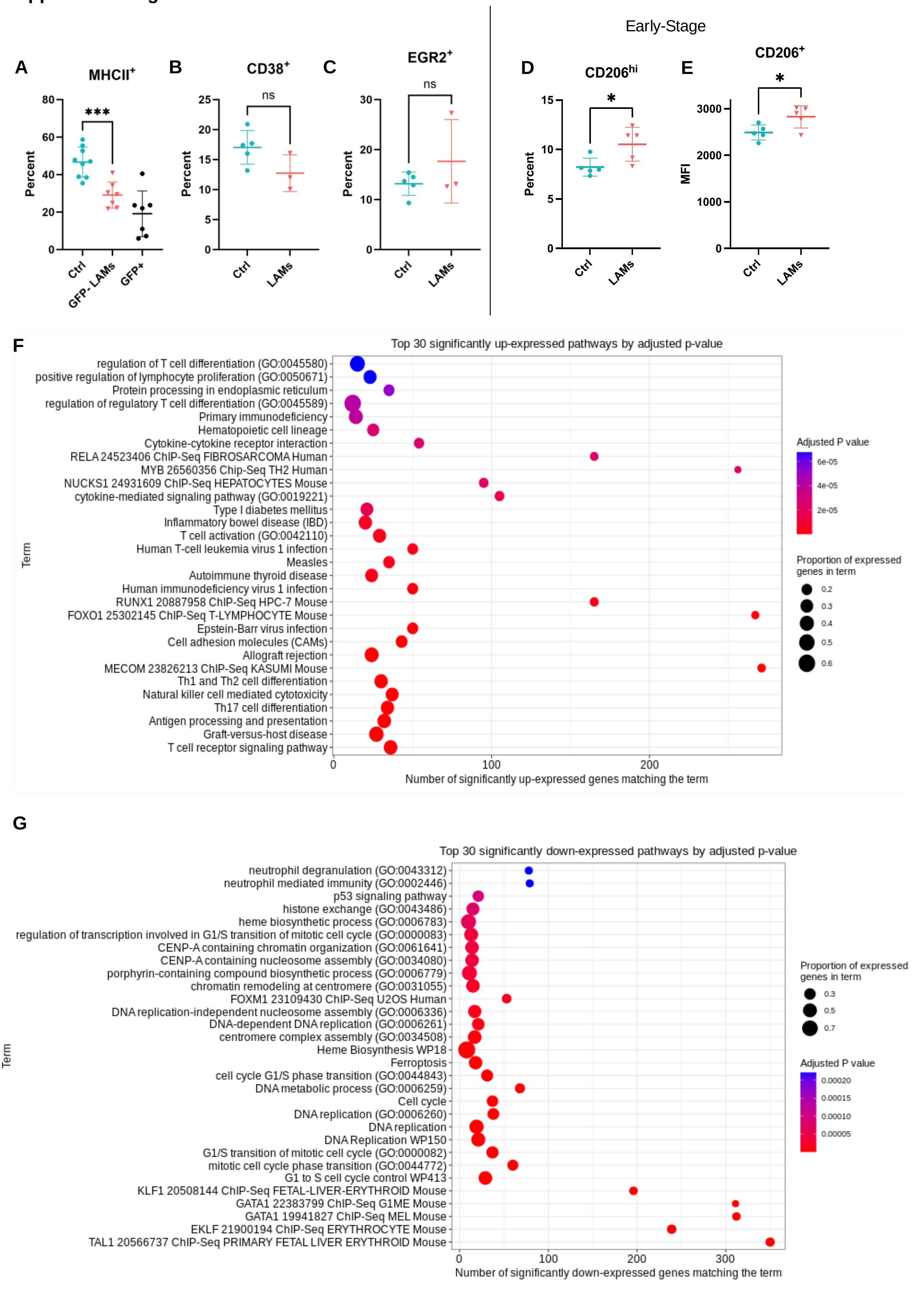

**Supplemental Fig. 2 continued**

**H**

**Enriched in TAMs**

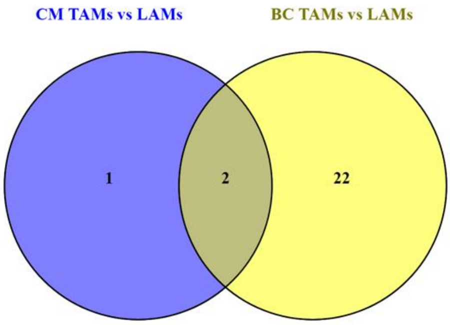


**Enriched in TAMs**

Both

BC

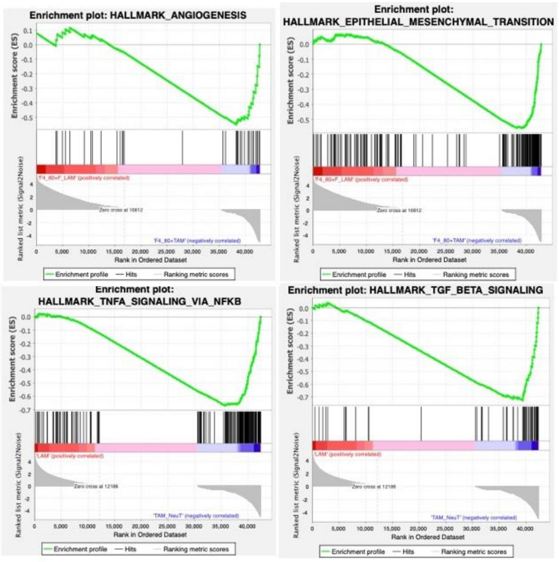

#### Supplemental Fig. 3

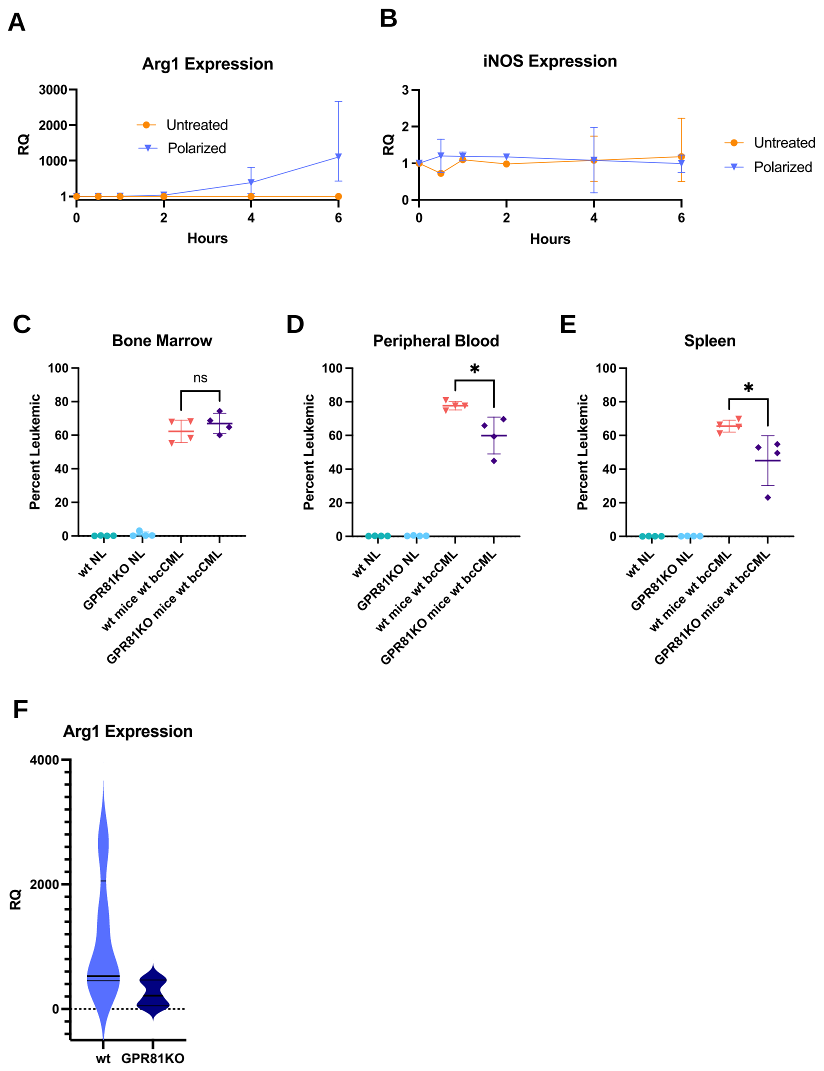

### Supplemental Fig. 4

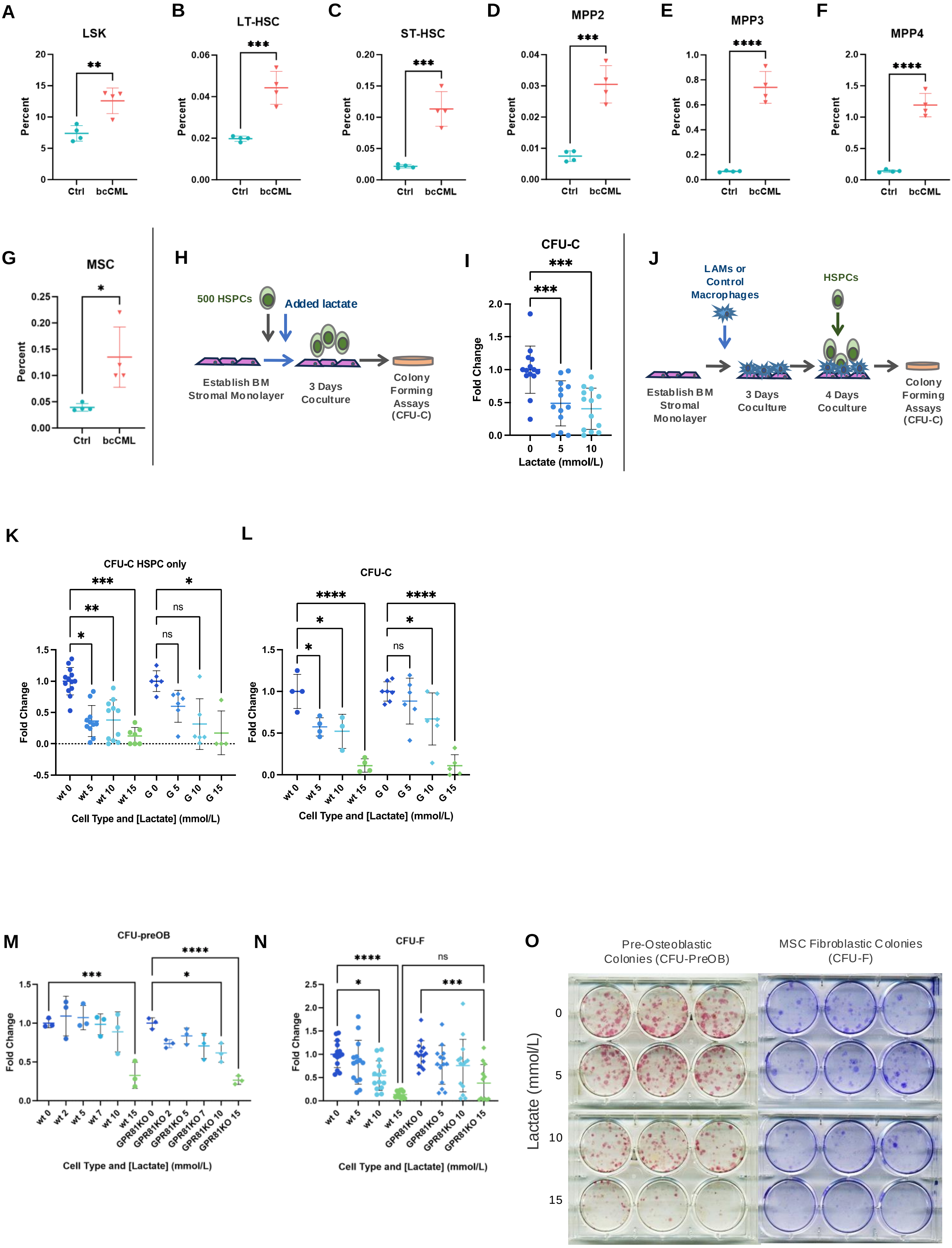

#### Supplemental Fig. 5

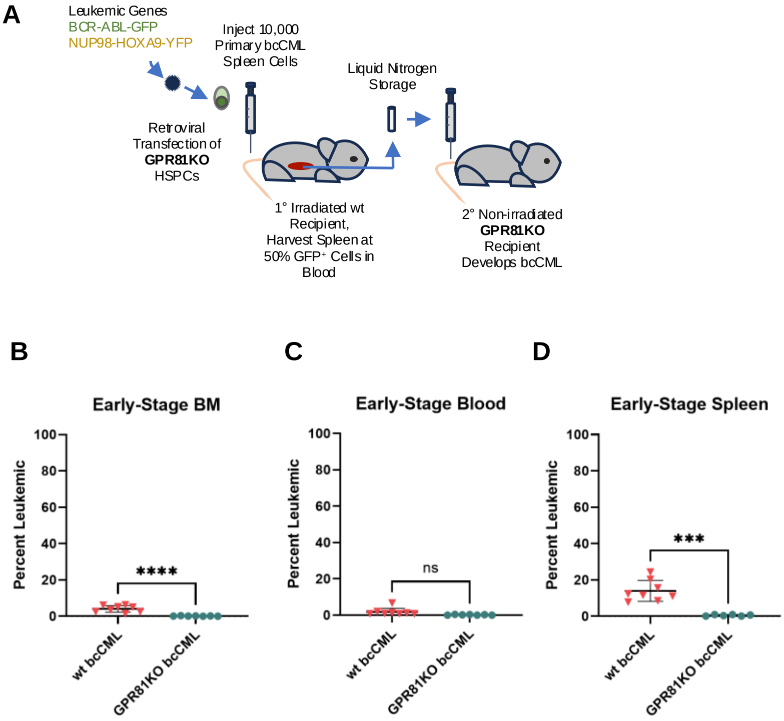
