## Supplemental Figure legends, methods, and tables for "Elevated Lactate in the AML Bone Marrow Microenvironment Polarizes Leukemia-Associated Macrophages via GPR81 Signaling"

**Fig. S1)** (**A**) Graphical depiction of bcCML cell generation and disease initiation. (**B**) Heatmap showing the relative abundance of detectable metabolites in bcCML bone marrow extracellular fluid, or healthy nonleukemic controls (n = 4, in triplicates). See **Table S2** for a list of significantly altered metabolites. (**C**) Mass spectrometry (MS) peak intensity of lactate measurements in human and murine BM extracellular fluid (n = 4 in triplicates).

**Fig. S2)** (**A-C**) BM macrophages at late-stage bcCML (55-70% leukemia cells in BM): Non-leukemia-derived LAMs ($GFP^-$ LAMs), leukemia-derived (GFP+). Macrophages from healthy nonleukemic controls (Ctrl). Percent positive for MHCII (n = 7-9) (**A**), one-way ANOVA was used, and for CD38 (n = 3-5) (**B**) or EGR2 (n = 3-5) (**C**), student's *t* test was used. (**D-E**) BM macrophages at early-stage disease (5% leukemia in the BM). Frequency of $CD206^{hi}$ (**D**) and expression of CD206 (n = 5) (**E**), student's *t* test was used. (**F-G**) Differentially expressed gene ontology (GO) elements from RNAseq analysis of LAMs compared to nonleukemic control macrophages: significantly up-expressed (**F**) or down-expressed (**G**) pathways identified by EnrichR analysis (n = 6). (**H**) Gene set enrichment analysis (GSEA), hallmark gene sets enriched in TAMs compared to LAMs. Venn diagram of number of significantly upregulated gene sets, and enrichment plots of top representative gene sets, determined by an FDR q-value of < 0.25. Data are represented with mean ± SD. Ns, not significant; *, P < 0.05; ***, P < 0.001. BC, breast cancer; CM, colorectal metastasis; Ctrl, nonleukemic control; LAMs, leukemia-associated macrophages; TAMs, tumor-associated macrophages.

**Fig. S3)** (**A-B**) Relative quantification (RQ) of *Arg1* (A) or *iNOS* (B) transcripts in untreated or polarized BMDMs for 0-6 hours (n = 5, in duplicates), error bars indicate mean with range. (**C-E**) Percent leukemic cells in the bone marrow (**C**), peripheral blood (**D**), or spleen (**E**) of wt or

GPR81KO mice with late-stage bcCML. (**F**) Relative quantification (RQ) of *Arg1* expression by wt or GPR81KO BMDMs polarized for 6 hours (n = 5, in duplicates), violin plot, lines represent interquartile range (IQR).

**Fig. S4)** (**A-G**) Hematopoietic progenitors' frequency of nonleukemic live cells in BM of late-stage bcCML mice or nonleukemic controls (Ctrl): Hematopoietic stem and progenitor cells (HSPCs/LSK) (**A**), long-term hematopoietic stem cells (LT-HSC) (**B**), short-term HSC (ST-HSC) (**C**), MPP subsets MPP2 (**D**)**,** MPP3 (**E**), and MPP4 (**F**) (n = 4), and frequency of stromal supportive mesenchymal stem cells (MSC) in bone (**G**), student's *t* test was used. (**H-I**) HSPCs cocultured on a BM stromal monolayer for 72 hours with or without lactate treatment: graphic of experimental procedure (**H**) and fold change CFU-Cs relative to the 0 mmol/L lactate control group (**I**) (n = 12-14), one-way ANOVA was used. (**J**) HSPCs were cocultured with a BM stromal monolayer and either 60,000 added LAMs or healthy control (Ctrl) macrophages for four days, graphical depiction of procedure. (**K-L**) CFU-Cs of wt or GPR81KO (G) lactate-treated HSPCs cultured alone (**K**) (n = 4-13) or cocultured with a stromal monolayer (**L**) (n = 3-7), relative to the 0 mmol/L lactate control group of the same genetic background. (**M-O**) wt or GPR81KO MSCs were treated with lactate while differentiated to preosteoblastic colonies (CFU-preOB) (n=4-13) (**M**), or cultured to fibroblastic colonies (CFU-F) (n=3-7) (**N**), representative images (**O**). Fold change CFU is shown relative to the 0 mmol/L lactate control of the same genetic background, one-way ANOVAs were used. Data are represented as mean ± SD. ns, not significant; *, $P < 0.05$; **, $P < 0.01$; ***, $P < 0.001$; ****, $P < 0.0001$. BM, bone marrow; CFU-C, colony forming unit cells; CFU-F, fibroblastic colony forming units; LAM, leukemia-associated macrophage; HSPC, hematopoietic stem and progenitor cell; LSK, lin-/ cKit+/Sca1+ HSPCs; MPP, multipotent progenitor; MSC, mesenchymal stem cells; pre-OB, preosteoblastic.

**Fig. S5)** (**A**) Graphical depiction of the procedure to produce GPR81KO bcCML cells, and initiation in GPR81KO mice (GPR81 DKO). (**B-D**) Leukemic burden in the BM (**B**), peripheral blood (**C**), and spleen (**D**) of wt or GPR81KO bcCML by the timepoint of wt early-stage disease (~5% leukemia in the BM) (n = 5), significance levels were determined by unpaired student's *t* tests. Data are represented as mean ± SD. ns, not significant; ***, $P < 0.001$; ****, $P < 0.0001$.

#### Supplemental Methods

**Metabolomics by Liquid Chromatography Mass Spectrometry (LC-MS/MS)**

Sample dilutions were prepared by mixing 20 µl sample and 280 µl cold 80:20 high-performance liquid chromatography (HPLC)-grade MeOH:water (Omnisolv MeOH MX0488-1; Omnisolv water WX0004-1) and briefly vortexed. Diluted samples were centrifuged at 4°C, at 16000 x g for 10 min to pellet precipitate, and the resulting supernatants were transferred to new tubes. For amino acids, samples underwent carbobenzyloxy (CBZ) derivatization by mixing 90 µl of diluted sample, 5 µl trimethylamine (Sigma 471283), and 1 µl benzyl chloroformate (ThermoFisher Acros Organics 152940050). Tubes were incubated at room temperature for 5 min then centrifuged at 16000 xg for 10 min, and supernatants were transferred to HPLC vials. For amino acids, a Synergi 4 µm Hydro-RP 80A 50x2mm column (Phenomenex 00B-4375-B0) was utilized with the following LC parameters: autosampler temperature, 4 °C; injection volume, 10 µL; column temperature, 40 °C; and flow rate, 0.5 mL/min. The LC solvents used were: solvent A, 100% methanol; and solvent B, 10 mM tributylamine and 15 mM acetic acid in 97:3 (vol:vol) water:methanol. The gradient conditions used were (vol/vol): negative mode-t = 0, 85% B; t = 4, 3%B; t = 5, 3% B; t = 5.1, 85% B. For all non-amino acid metabolites, 100 µl of methanol extracts were cleared by centrifugation as above, transferred to HPLC vials, and 10 µL injected. These extracts were run over a Synergi 4 µm hydro-RP 80A 150x2mm HPLC column (Phenomenex (Torrance, CA, USA) 00F-4375-B0). The total run time was 20 min, with solvent A being 97:3 water/methanol with 10 mM tributylamine and 15 mM acetic acid; and solvent B being methanol. The gradient was 0 min, 0% B, flow rate 200 µL/min; 2.5

min, 0% B, flow rate 200 μL/min; 5 min, 20% B, flow rate 250 μL/min; 12 min, 95% B, flow rate 250 μL/min; 14.5 min, 95% B, flow rate 250 μL/min; 15 min, 0% B, flow rate 250 μL/min; 19 min, 0% B, flow rate 200 μL/min; 20 min, 0% B, flow rate 200 μL/min.

*Data Processing and Analyses*: Raw Xcalibur data was transformed to peak intensity (relative abundance) of known compounds using mzRock analysis software (https://code.google.com/archive/p/mzrock/). Compound prediction reports were used to manually filter any trace detection (use only those with aligned retention time, intensity >10000). MetaboAnalyst 5.0 software (RRID:SCR_015539) was used to process the peak intensity data, perform the statistical analysis, and generate heatmaps. Peak intensity values were uploaded. There were no missing values and no data was filtered. The data were not normalized. Orthogonal partial least squares – discriminant analysis (orthoPLS-DA) was performed to generate a scores plot. T tests were performed with false discovery rate (FDR) correction. Clustering/heatmaps were done using the Euclidean distance measure and ward.D clustering algorithm, then sample triplicates were presented as grouped together by clicking “do not cluster” on the heatmap analyses table.

**Murine Strains and Ethics**

All murine experiments were performed using male and female wild type C57BL/6J mice obtained from the Jackson Laboratory (RRID:IMSR_JAX:000664). Previous studies from our lab and others have shown no sex-dependent differences in the BMME in the number of HSCs, mesenchymal stem cells (MSCs), macrophages, endothelial cells, osteoblastic cells (OBs), or the immunohistochemistry involved. The male and female data were plotted together unless a sex-specific difference was noted. All mice were maintained by monogamous breeding and housed pathogen-free at the Vivarium at the University of Rochester School of Medicine and Dentistry.

*Gpr81*$^{-/-}$ mice, genetically modified to lack a functional gene for the receptor GPR81/HCAR1, previously characterized and used for published studies: They were kindly acquired via material transfer agreement from Dr. Vadivel Ganapathy, Ph.D. (professor and department chair of cell biology and biochemistry at Texas Tech University Health Sciences Center, USA), who obtained the mice from Dr. Stefan Offermanns (professor and director of the department of pharmacology at Max-Planck-Institute for Heart and Lung Research, Germany). Mice were on a B6 (>10 generation) background [1]. Dr. Offermanns originally procured the mice from the Texas Institute of Genomic Medicine (Houston, TX, USA) [2].

Preliminary studies were used to determine a sample size calculation to utilize minimal numbers of research animals. Subjects were excluded from count if they experienced health concerns for any reason other than the intended leukemia model. Experiments were limited to critical *in vivo* mechanisms that could not be assayed appropriately *in vitro*. Approval by the Institutional Animal Care and Use Committee (IACUC) and the Institutional Review Board (IRB) were acquired for this research. All facilities and animal care comply with federal and NIH policies, accredited by the Association for Assessment and Accreditation of Laboratory Animal Care International (AAALAC).

**Murine AML Model (bcCML)**

BcCML, which includes lentiviral insertion of both the BCR-ABL translocation conjugated to GFP, as well as Nup-Hox98 fusion conjugated to YFP, has been previously characterized [3-7]. Primary bcCML was initiated via tail vein injection of bcCML cells into 8–12-week-old mice, post 6.5 gray (Gy) sublethal irradiation. For comparative bcCML *in vivo* experiments, 10,000 wt or *Gpr81*$^{-/-}$ bcCML cells per 100 µL dose were injected into age- and sex-matched wt or *Gpr81*$^{-/-}$ mice.

**Whole Bone Marrow Isolation from Mice**

6-12-week-old mice were euthanized by carbon dioxide asphyxiation followed by cervical dislocation. Mice were briefly dipped before dissection into a container of 70% ethanol (200-proof ethanol (Avantor VWR (Radnor, PA, USA) 89125-174) diluted using distilled water ($dH_20$). The hindlimbs (tibias and femurs) were removed by dissection, and the muscles were removed from the outside of the bone by dissection followed by rubbing with a Kimtech Kimwipe (Kimberly-Clark (Irving, TX, USA) 34256). The bones were crushed in 1X PBS using a pestle and mortar to remove the BM in 5 mL of media, then filtered through a 40 μM cell strainer into a 50 mL conical tube to remove bone fragments. The cells were pelleted by centrifugation at 900 x g for five minutes and the supernatant was removed.

**Murine Bone Marrow Extracellular Fluid Collection**

Tibias and femurs were dissected and crushed in 1 mL of 1X PBS. This cellular suspension was then filtered through a 40 μm cell strainer into a 50 mL conical tube to remove bone fragments. The cells were pelleted by centrifugation at 900 x g for five minutes and the supernatant was removed into an Eppendorf tube, then immediately stored at -80C until use.

**Flow cytometry and fluorescence-activated cell sorting (FACS)**

Bone marrow samples underwent red blood cell lysis prior to flow cytometric analyses**:** Whole bone marrow freshly isolated from mice was resuspended in red blood cell lysis buffer (156 mM $NH_4CL$ (Sigma 213330-25G), 127 μM EDTA (Sigma E4884-100G), and 12 mM $NaHCO_3$ (Sigma S6014-25G) in $dH_20$ (all $dH_20$ for cellular or molecular work was purified by use of a Barnstead Nanopure system (ThermoFisher Scientific)) for five minutes to lyse the red blood cells before flow cytometry analysis. Cells are resuspended in a greater amount of 1X PBS then centrifuged to the pellet, and the supernatant is removed.

Samples were resuspended in 1X phosphate-buffered saline (PBS) (Corning 21-040-CV) with added 2% heat-treated fetal bovine serum (htFBS) (Gibco 26140079, heat-treated at 56°C for 30 min). All antibodies for flow cytometry and cell sorting were obtained commercially. See **Supplemental Methods Table 1 (Table SM1)** for a list of flow cytometry antibodies and **Supplemental Methods Table 2 (Table SM2)** for markers and gating strategies used. Fluorescence minus one (FMO) were prepared using cells, and positive controls were prepared using UltraComp eBeads compensation beads (ThermoFisher Invitrogen 01-2222-42), in each experiment to ensure accurate staining and appropriate gating. Only the live, single cells were considered in analyses, determined by side scatter and forward scatter and 4',6-diamidino-2-phenylindole (DAPI) live/dead nuclear stain.

**Simplified Presentation of Incredibly Complex Evaluations (SPICE)**

The SPICE analysis was performed using SPICE 6 software (RRID:SCR_016603) publicly available through the NIH NIAID site.

**RNA sequencing (RNAseq)**

Demultiplexing, quality control, alignment, and analysis methods: Raw reads generated from the Illumina basecalls were demultiplexed using bcl2fastq version 2.19.1. Quality filtering and adapter removal are performed using FastP version 0.23.1 (RRID:SCR_016962) with the following parameters: "--length_required 35 --cut_front_window_size 1 --cut_front_mean_quality 13 --cut_front --cut_tail_window_size 1 --cut_tail_mean_quality 13 --cut_tail -y –r" [8]. Processed/cleaned reads were then mapped to the GRCm39/gencode M31 reference using STAR_2.7.9a with the following parameters: "—twopass Mode Basic --runMode alignReads --outSAMtype BAM Unsorted – outSAMstrandField intronMotif --outFilterIntronMotifs RemoveNoncanonical –outReadsUnmapped Fastx" [9,10]. Genelevel read quantification was derived using the subread-2.0.1 package (featureCounts, RRID:SCR_012919) with a GTF annotation file GRCm39/gencode M31, and the following parameters for stranded RNA libraries "-s 2 -t exon -g gene_name" [11]. Differential expression analysis was performed using DESeq2-1.34.0 with a P-value threshold of 0.05 within R version 3.5.1 (https://www.R-project.org/) [12]. A PCA plot was created within R using the pcaExplorer to measure sample expression variance [13]. Heatmaps were generated using the pheatmap package (RRID:SCR_016418) using rLog transformed expression values [14]. Gene ontology analyses were performed using the EnrichR package (RRID:SCR_001575) [15-17] Volcano plots and dot plots were created using ggplot2 (RRID:SCR_014601) [18].

**Comparative Transcriptomics**

Murine TAM RNA sequencing datasets were accessed via GEO Accession viewer IDs GSE206211 and GSE126268. In our study, we analyzed both male and female F4/80$^+$ LAMs. The controls that were used for the LAM analysis were F4/80+ macrophages from healthy wt mice of the same age, and sex, and from the same background (C57BL/6J). The GSE126268 Tuit et al. study analyzed female F4/80$^+$ tumor-associated macrophages from a NeuT murine breast cancer model, and for controls used healthy F4/80$^+$ wt littermates (from the same BALB/c background). The GSE206211 Qiao et al. study analyzed female F4/80$^+$ tumor-associated macrophages from a colorectal liver metastasis murine model, and the controls used were macrophages from livers with or without metastasis.

*Alignment, and Analysis*:

Raw fastq files were downloaded from GEO using the SRA run selector. Processed/cleaned reads were then mapped to the GRCm39/gencode M31 reference using STAR_2.7.9a with the following parameters: "—twopass Mode Basic --runMode alignReads --outSAMtype BAM Unsorted –outSAMstrandField intronMotif --outFilterIntronMotifs RemoveNoncanonical –outReadsUnmapped Fastx" [10,19]. Gene level read quantification was derived using the subread-2.0.1 package (featureCounts) with a GTF annotation file (GRCm39/gencode M31) and the following parameters for stranded RNA libraries "-s 2 -t exon -g gene_name" [10,20]. Differential expression analysis was performed using DESeq2-1.34.0 with a P-value threshold of 0.05 within R version 4.0.2 (https://www.R-project.org/) [12]. A PCA plot was created within R using pcaExplorer c2.14.2 to measure sample expression variance [13]. Heatmaps were generated using the pheatmap 1.0.12 package with rLog transformed expression values [14]. Gene ontology analyses were performed using the EnrichR package [15,21,22]. Volcano plots and dot plots were created using ggplot2 [18].

Pathway analysis: To compare cancer-associated macrophage types, the Enrichr results were used to create the Venn diagrams by use of the VENNY 2.1 online visualizer (BioinfoGP, CNB-CSIC) [23]. To perform an analysis consistent to all types of cancer-associated macrophages, all data was normalized to our healthy BM macrophage female controls, and only the female LAMs were considered.

*Gene Set Enrichment Analysis (GSEA)*: The GSEA analysis was performed using the DESeq2 (RRID:SCR_000154) normalized read counts and processed with GSEA_4.3.2. Significance was defined as FDR is < 0.25. As the TAM studies used female macrophages, we only considered our female LAM samples. For consistency, all groups were compared to our nonleukemic macrophage controls' RNA expression as a baseline. Here, we queried hallmark genes, which are associated with well-defined biological states/pathways that have homology in mice and humans.

**Quantitative real-time polymerase chain reaction (qRT-PCR)**

Cells were grown to 70% confluency in corresponding media in tissue culture-treated 12-well plates. Media was changed to serum-free media for 12-24 hours before lactate treatment. Lactate was added to wells at 10 mmol/L and cells were cultured for 0-6 hours, and 5 ng/mL of IL4 and IL13 was also added to stimulate polarization. Cells were then removed from the well by treatment with 0.25% trypsin-EDTA for three minutes then by the additional use of a cell-scraper for macrophage cultures. All cells were collected in an Eppendorf tube and centrifuged at 3000 x g for 3 minutes to pellet the cells. The supernatant was removed, and cells were resuspended in RLT lysis buffer (Qiagen), then stored at -20C until RNA extraction by RNeasy Plus Mini Kit (Qiagen 74134). The cDNA libraries were then prepared using the High-Capacity cDNA Reverse Transcription kit (Applied Biosystems, ThermoFisher 4368814). The TaqMan Gene Expression Master Mix (ThermoFisher 4369016) was used for qRT-PCR along with the following TaqMan Gene Expression Assays (FAM) (ThermoFisher 4331182): Arg1 mouse (Mm00475988_m1), iNOS mouse (Mm00440502_m1), and beta-actin mouse (Mm04394036_g1). The assay was run on the QuantStudio 12KFlex Real-Time PCR System at the UR Genomics Research Center. The relative quantification (RQ) of mRNA expression was calculated using the $2^{-(\Delta\Delta Ct)}$ method (Ct = cycle threshold), sample calculations were normalized to beta-actin.

**Murine Complete Blood Counts (CBCs)**

Blood samples were collected from mice by submandibular bleeds into BD Microtainer tubes with K2E ($K_2EDTA$) (Becton Dickinson (BD) (Franklin Lakes, NJ, USA) Biosciences 365974). CBCs were performed on 10 µl of blood using a scil Vet abc Plus hematology analyzer (Antech Diagnostics (Fountain Valley, CA, USA)).

**Colony forming unit (CFU-C) assays**

*LSK with Stromal Monolayer*: Whole BM freshly isolated from mice was plated on six-well cell culture-treated dishes at $4x10^6$ cells per well in complete MEM α without ascorbic acid. Cells were grown overnight, then non-adherent cells were removed from the dish and fresh media was added. Cells were grown to confluency, with media aspirated and fresh media added every 3 - 4 days. LSKs were sorted via FACS and added on day 14 in LSK media. The cocultures were incubated for 72 hours and then plated for colony-forming units CFU-Cs.

*LSK Only*: These were cultured for 72 hours in a sterile 24-well suspension culture plate (Greiner Bio-One (Kremsmünster, Austria) CELLSTAR 622102) in LSK media with varying lactate treatments, then plated for CFU-Cs.

*With Added LAMs*: 60,000 macrophages were obtained by FACS from the BM of either leukemic mice or non-leukemic control mice and plated into the wells to grow for 4 days. Then, LSKs were sorted from freshly isolated mouse BM, and 500 cells per well were added to the stromal coculture support layer. These were cultured for four days in LSK media with or without lactate added, then plated for CFU-C.

**Colony Forming Unit-Osteoblast (CFU-OB) Assay**

Whole BM from mice were plated at $4 \times 10^6$ cells per well in tissue culture-treated 6-well plates, or at $2 \times 10^6$ cells per well in 12-well plates, in ascorbic acid-free MEM α at 37°C, 5% CO2. After attachment overnight, media with nonadherent cells was removed, and fresh media was added. On day four, the media was changed to mineralizing media (complete MEM α without ascorbic acid was used as a media base, with added 0.05 mg/mL L-ascorbic acid (Krackeler Scientific (Albany, NY, USA) 45-A4544) and 10mM glycerol 2-phosphate disodium salt hydrate (Krackeler Scientific G9422) then sterile-filtered (NEST 343011). Cells were treated with 0, 5, 10, or 15 mmol/L of excess lactate starting on day 4, and incubated for 14 days, with media changed every three days. The attached cells were fixed with 10% neutral buffered formalin for 20 minutes and then rinsed with $dH_20$. Triplicate wells were assayed for alkaline phosphatase activity, indicating pre-osteoblasts (Pre-OB), by staining for 45 minutes with a solution of 100 mM Tris-HCl, 0.005% weight/volume naphthol AS MX-PO4, and 0.03% weight/volume red violet LB salt. To assay for silver nitrate-positive colonies, cultures were grown until day 28, and then Von Kossa staining was done by staining for alkaline phosphatase followed by three rinses with $dH_20$ and then 2.5% silver nitrate solution for 30 minutes. After staining, wells were then rinsed with $dH_20$ and allowed to dry. Plates were imaged using a high-definition scanner (Canon (Tokyo, Japan)), and then images were analyzed for area by color thresholding using ImageJ image analysis software (NIH, RRID:SCR_003070).

##### Colony Forming Unit-Fibroblast (CFU-F) and MSC Proliferation Assays

Whole BM from mice were plated at P0 at 2 x $10^6$ cells per well in tissue culture-treated 12-well plates in ascorbic acid-free MEM α at 37°C, 5% CO2. Cells were treated with 0, 5, 10, or 15 mmol/L of excess lactate starting on day 4, and incubated for 10-14 days, with media changed every three days. The attached cells were fixed with 10% neutral buffered formalin for 20 minutes, then rinsed with $dH_20$. Wells were stained with a 1:1 mix of 1% crystal violet dissolved in 100% methanol and $dH_20$ for 30 minutes, rinsed with $dH_20$ three times, and left in water overnight. To quantify crystal violet staining, the water was removed and rinsed, then the crystal violet stain was solubilized in 300 µl of 10% acetic acid in dH20 for 15 minutes, then diluted 1:2 in $dH_20$, and the absorbance was read at 590 nm on the microplate reader.

##### Leukemia Stem Cell (LSC) Repopulation Assay

Doubly-transfected ($GFP^+$, $YFP^+$) wt or *Gpr81*$^{-/-}$ primary bcCML cells were sorted from spleen by FACS. Cells were plated in MethoCult M3434 as described in CFU-C assays. After 14 days, four well-established colonies were picked, resuspended in 200 µl of media, then replated.

##### Scientific Rigor

For randomization of non-intended influential variables, experiments were repeated three times with separate starting dates, and the data was compiled. Counts were blinded to the group to inhibit bias. Power caclulation analyses were performed prior to repeats of the experiment.

#### Supplemental Methods References

14. Kolde R. Package ‘pheatmap’. Version 1.0.12.

15. Chen EY, Tan CM, Kou Y, et al. Enrichr: interactive and collaborative HTML5 gene list enrichment analysis tool. *BMC Bioinformatics*. 2013;14(1):128.

16. Kuleshov MV, Jones MR, Rouillard AD, et al. Enrichr: a comprehensive gene set enrichment analysis web server 2016 update. *Nucleic Acids Research*. 2016;44(W1):W90-W97.

17. Xie Z, Bailey A, Kuleshov MV, et al. Gene Set Knowledge Discovery with Enrichr. *Current Protocols*. 2021;1(3):e90.

18. Wickham H. ggplot2: Elegant Graphics for Data Analysis: Springer-Verlag New York; 2016.

### Supplemental Methods Tables

| Marker | Antibody to | Fluor/Conjugate | Company | Brand | Product # |
|---|---|---|---|---|---|
| live/dead cell | n/a | DAPI | ThermoFisher Scientific | Invitrogen | D21490 |
| monocytes, neutrophils | Ly-6C | APC-eFluor 780 | ThermoFisher Scientific | Invitrogen | 47-5932-82 |
| monocytes, granulocytes, neutrophils | Ly-6G | Brilliant Violet 786 | BD Biosciences | OptiBuild | 740953 |
| hematopoietic cells | CD45 | APC | BioLegend (San Diego, CA, USA) | | 103112 |
| murine macrophages | F4/80 | Brilliant Violet 650 | BioLegend | | 123149 |
| murine macrophages | F4/80 | PE | ThermoFisher Scientific | Invitrogen | 12-4801-82 |
| "classic" activated macrophages | CD38 | Alexa Fluor700 | ThermoFisher Scientific | Invitrogen eBioscience | 56-0381-82 |
| "suppressive" alternative macrophages | EGR2 | PE | Miltenyi | | 130-114-363 |
| "classic" activated macrophages | MHCII | PE-Cyanine5 | BioLegend | | 107611 |
| "suppressive" alternative macrophages | CD206 | PE-Cyanine7 | ThermoFisher Scientific | Invitrogen | 25-2061-82 |
| lineage | CD3e | Biotin | ThermoFisher Scientific | Invitrogen | 13-0031-82 |
| lineage | Ly-6G/Ly-6C (Gr-1) | Biotin | ThermoFisher Scientific | Invitrogen | 13-5931-82 |
| lineage | TER-119 | Biotin | ThermoFisher Scientific | Invitrogen | 50-120-40 |
| lineage | B220 | Biotin | ThermoFisher Scientific | Invitrogen | 13-0452-82 |
| lineage, streptavidin 2° | biotin | Brilliant Violet 650 | BioLegend | | 405232 |
| hematopoietic stem cells | stem cell antigen-1 (Sca-1) | PerCP-Cyanine5.5 | ThermoFisher Scientific | Invitrogen | 45-5981-82 |
| hematopoietic progenitor cells | CD117 (c-Kit) | PE-Cyanine5 | ThermoFisher Scientific | Invitrogen eBioscience | 15-1171-83 |
| hematopoietic stem/progenitor cells | CD135 (Flt3) | PE | ThermoFisher Scientific | Invitrogen eBioscience | 12-1351-82 |
| MPPs, lymphocytes and monocytes | CD48 | APC-Cyanine7 | BioLegend | | 103432 |
| hematopoietic stem cells | CD150 | Brilliant Violet 785 | BioLegend | | 115937 |
| hematopoietic progenitor cells | CD117 (c-Kit) | PE-Cyanine5 | ThermoFisher Scientific | Invitrogen | 15-1171-83 |
| erythroid cells | TER-119 | FITC | BD Biosciences | Pharmingen | 557915 |
| hematopoietic cells | CD45 | APC-Cyanine7 | BD Biosciences | Pharmingen | 561037 |

Table SM1: List of α-Murine Antibodies Used for Flow Cytometry

| Cell Type | Gating Strategy |
|---|---|
| Macrophage | $ly6c^-$, $ly6g^-$, $CD45^+$, $F4/80^+$ |
| HSPC (LSK) | lineage $(lin)^-$ (lin = B220, Ter119, Ly-6C/Ly-6G (Gr-1), and CD3e), $c\text{-}Kit^+$, $Sca\text{-}1^+$ |
| LT-HSC | LSK, $Flt3^-$, $CD150^+$, $CD48^-$ |
| ST-HSC | LSK, $Flt3^-$, $CD150^-$, $CD48^-$ |
| MPP2 | LSK, $Flt3^-$, $CD150^+$, $CD48^+$ |
| MPP3 | LSK, $Flt3^-$, $CD150^-$, $CD48^+$ |
| MPP4 | LSK, $Flt3^+$, $CD150^-$, $CD48^+$ |

Table SM2: Markers and Gating Strategies for Flow Cytometry Populations

### Supplemental Tables

| Group | Sex | Age | Mutations at Diagnosis from BM Biopsy |
|---|---|---|---|
| AML | M | 31 | IDH1, NRAS, RUNX1 |
| AML | M | 33 | ETV6, NRSAS, RUNX1 |
| AML | F | 37 | Biallelic CEBPα mutated; no NGS done; karyotype normal |
| AML | F | 67 | Inv 16; no mutations |
| Control | M | 30 | |
| Control | M | 30 | |
| Control | F | 35 | |
| Control | F | 67 | |

Table S1: Human BM Serum Sample Metrics: Age, Sex, and Mutations

| | t.stat | p.value | -log10(p) | FDR |
|---|---|---|---|---|
| **ribose-P** | 29.585 | 3.28E-19 | 18.485 | 1.34E-17 |
| **sedoheptulose-P** | 11.411 | 1.04E-10 | 9.9839 | 2.13E-09 |
| **PEP** | 7.862 | 7.87E-08 | 7.104 | 9.12E-07 |
| **DHAP** | 7.8035 | 8.90E-08 | 7.0507 | 9.12E-07 |
| **tryptophan** | 6.6503 | 1.10E-06 | 5.9599 | 8.99E-06 |
| **hexose-P** | 6.2652 | 2.64E-06 | 5.5789 | 1.80E-05 |
| **F6P** | 6.0841 | 4.01E-06 | 5.3972 | 2.07E-05 |
| **proline** | 6.0812 | 4.03E-06 | 5.3942 | 2.07E-05 |
| **G6P** | 5.7494 | 8.77E-06 | 5.057 | 4.00E-05 |
| **GMP** | 5.6134 | 1.21E-05 | 4.9173 | 4.96E-05 |
| **glutamate** | 5.4096 | 1.97E-05 | 4.7065 | 7.33E-05 |
| **IMP** | 5.3028 | 2.54E-05 | 4.5954 | 8.67E-05 |
| **malate** | 5.2374 | 2.97E-05 | 4.5271 | 9.37E-05 |
| **gluconate** | 4.9808 | 5.52E-05 | 4.2581 | 0.00015891 |
| **AMP** | 4.9593 | 5.81E-05 | 4.2355 | 0.00015891 |
| **G3P** | -4.8982 | 6.74E-05 | 4.1711 | 0.00017281 |
| **phenylalanine** | 4.7998 | 8.56E-05 | 4.0673 | 0.00020654 |
| **alanine** | 4.7734 | 9.13E-05 | 4.0394 | 0.000208 |
| **leucine/isoleucine** | 4.629 | 0.00012984 | 3.8866 | 0.00027863 |
| **NAG** | 4.6102 | 0.00013592 | 3.8667 | 0.00027863 |
| **UDP-D-glucose** | 4.3548 | 0.00025358 | 3.5959 | 0.00049508 |
| **ornithine** | 4.3277 | 0.000271 | 3.567 | 0.00050505 |
| **tyrosine** | 4.3009 | 0.00028931 | 3.5386 | 0.00051573 |
| **asparagine** | 4.1152 | 0.00045539 | 3.3416 | 0.00077796 |
| **methionine** | 3.4358 | 0.0023603 | 2.627 | 0.0038709 |
| **glutathione** | -3.3772 | 0.002715 | 2.5662 | 0.0042813 |
| **lactate** | 3.1531 | 0.0046143 | 2.3359 | 0.0070069 |
| **lysine** | 2.9497 | 0.0074084 | 2.1303 | 0.010848 |
| **histidine** | 2.8375 | 0.009581 | 2.0186 | 0.013546 |
| **aspartate** | -2.741 | 0.011925 | 1.9235 | 0.016298 |
| **threonine** | 2.6213 | 0.015591 | 1.8071 | 0.02062 |
| **pantothenate** | 2.5115 | 0.019863 | 1.702 | 0.025377 |
| **arginine** | -2.4987 | 0.020425 | 1.6898 | 0.025377 |
| **serine** | 2.2731 | 0.033137 | 1.4797 | 0.03996 |

Table S2: Altered Extracellular Metabolites in Murine bcCML Bone Marrow

| Common Downregulated Elements | | |
|---|---|---|
| **LAMs, CM and BC TAMs** | **LAMs and CM TAMs** | **LAMs and BC TAMs** |
| DNA replication initiation (GO:0006270) | histone exchange (GO:0043486) | nuclear DNA replication (GO:0033260) |
| DNA-dependent DNA replication (GO:0006261) | DNA replication-independent nucleosome assembly (GO:000633 | regulation of transcription involved in G1/S transition of mitotic cell cycle (GO:0000083) |
| DNA replication (GO:0006260) | positive regulation of cell cycle G2/M phase transition (GO:1902 | DNA replication checkpoint signaling (GO:0000076) |
| mitotic chromosome condensation (GO:0007076) | iron-sulfur cluster assembly (GO:0016226) | chromosome condensation (GO:0030261) |
| mitotic sister chromatid segregation (GO:0000070) | regulation of cyclin-dependent protein kinase activity (GO:19040 | organonitrogen compound biosynthetic process (GO:1901566) |
| mitotic cell cycle phase transition (GO:0044772) | DNA strand elongation (GO:0022616) | protein-DNA complex assembly (GO:0065004) |
| heme biosynthetic process (GO:0006783) | base-excision repair (GO:0006284) | proteasome-mediated ubiquitin-dependent protein catabolic process (GO:0043161) |
| centromere complex assembly (GO:0034508) | DNA repair (GO:0006281) | DNA damage response, signal transduction by p53 class mediator (GO:0030330) |
| DNA metabolic process (GO:0006259) | translesion synthesis (GO:0019985) | erythrocyte differentiation (GO:0030218) |
| porphyrin-containing compound biosynthetic process (GO:0006779) | Mismatch repair | DNA unwinding involved in DNA replication (GO:0006268) |
| DNA strand elongation involved in DNA replication (GO:0006271) | | DNA damage response, signal transduction by p53 class mediator resulting in cell cycle arrest (GO:0006977) |
| G1/S transition of mitotic cell cycle (GO:0000082) | | cellular macromolecule biosynthetic process (GO:0034645) |
| chromatin remodeling at centromere (GO:0031055) | | regulation of ubiquitin protein ligase activity (GO:1904666) |
| microtubule cytoskeleton organization involved in mitosis (GO:1902850) | | kinetochore assembly (GO:0051382) |
| histone exchange (GO:0043486) | | mitotic DNA damage checkpoint signaling (GO:0044773) |
| CENP-A containing chromatin organization (GO:0061641) | | Ferroptosis |
| CENP-A containing nucleosome assembly (GO:0034080) | | p53 signaling pathway |
| DNA replication-independent nucleosome assembly (GO:0006336) | | Progesterone-mediated oocyte maturation |
| regulation of mitotic cell cycle phase transition (GO:1901990) | | Oocyte meiosis |
| regulation of G2/M transition of mitotic cell cycle (GO:0010389) | | GATA1 19941827 ChIP-Seq MEL Mouse |
| cell cycle G2/M phase transition (GO:0044839) | | GATA1 22383799 ChIP-Seq G1ME Mouse |
| regulation of cell cycle G2/M phase transition (GO:1902749) | | AR 21909140 ChIP-Seq LNCAP Human |
| positive regulation of mitotic cell cycle phase transition (GO:1901992) | | MECOM 23826213 ChIP-Seq KASUMI Mouse |
| G2/M transition of mitotic cell cycle (GO:0000086) | | LMO2 20887958 ChIP-Seq HPC-7 Mouse |
| metallo-sulfur cluster assembly (GO:0031163) | | CREM 20920259 ChIP-Seq GC1-SPG Mouse |
| double-strand break repair via homologous recombination (GO:0000724) | | GATA2 22383799 ChIP-Seq G1ME Mouse |
| positive regulation of cell cycle G2/M phase transition (GO:1902751) | | E2F4 21247883 ChIP-Seq LYMPHOBLASTOID Human |
| mitotic spindle organization (GO:0007052) | | MYB 21317192 ChIP-Seq ERMYB Mouse |
| double-strand break repair via break-induced replication (GO:0000727) | | MYCN 18555785 ChIP-Seq MESCs Mouse |
| mitotic G1 DNA damage checkpoint signaling (GO:0031571) | | p53 signaling WP2902 |
| iron-sulfur cluster assembly (GO:0016226) | | |
| regulation of cyclin-dependent protein kinase activity (GO:1904029) | | |
| DNA integrity checkpoint signaling (GO:0031570) | | |
| sister chromatid segregation (GO:0000819) | | |
| kinetochore organization (GO:0051383) | | |
| mitotic DNA replication (GO:1902969) | | |
| DNA strand elongation (GO:0022616) | | |
| base-excision repair (GO:0006284) | | |
| DNA repair (GO:0006281) | | |
| pre-replicative complex assembly involved in nuclear cell cycle DNA replication (GO:0006267) | | |
| pre-replicative complex assembly (GO:0036388) | | |
| ubiquitin-dependent protein catabolic process (GO:0006511) | | |
| translesion synthesis (GO:0019985) | | |
| regulation of mitotic metaphase/anaphase transition (GO:0030071) | | |
| DNA replication | | |
| Cell cycle | | |
| Mismatch repair | | |
| TAL1 20566737 ChIP-Seq PRIMARY FETAL LIVER ERYTHROID Mouse | | |
| EKLF 21900194 ChIP-Seq ERYTHROCYTE Mouse | | |
| FOXM1 23109430 ChIP-Seq U2OS Human | | |
| FOXM1 25889361 ChIP-Seq OE33 AND U2OS Human | | |
| E2F7 22180533 ChIP-Seq HELA Human | | |
| G1 to S cell cycle control WP413 | | |
| DNA Replication WP150 | | |
| Heme Biosynthesis WP18 | | |

Table S3: Common Downregulated Elements in LAMs and TAMs

| Gene sets Enriched in LAM vs. CM TAMs | | Gene sets Enriched in LAM vs. BC TAMs | |
|---|---|---|---|
| **GS DETAILS** | **FDR q-val** | **GS DETAILS** | **FDR q-val** |
| HALLMARK_E2F_TARGETS | 0 | HALLMARK_MYC_TARGETS_V1 | 0 |
| HALLMARK_OXIDATIVE_PHOSPHORYLATION | 0 | HALLMARK_E2F_TARGETS | 0 |
| HALLMARK_MYC_TARGETS_V1 | 0 | HALLMARK_OXIDATIVE_PHOSPHORYLATION | 0 |
| HALLMARK_DNA_REPAIR | 0 | HALLMARK_MYC_TARGETS_V2 | 0 |
| HALLMARK_MYC_TARGETS_V2 | 0 | HALLMARK_DNA_REPAIR | 0 |
| HALLMARK_MTORC1_SIGNALING | 0 | HALLMARK_G2M_CHECKPOINT | 0 |
| HALLMARK_G2M_CHECKPOINT | 0 | HALLMARK_MTORC1_SIGNALING | 0.001 |
| HALLMARK_HEME_METABOLISM | 0 | HALLMARK_FATTY_ACID_METABOLISM | 0.001 |
| HALLMARK_UNFOLDED_PROTEIN_RESPONSE | 0 | HALLMARK_REACTIVE_OXIGEN_SPECIES_PATHWAY | 0.003 |
| HALLMARK_FATTY_ACID_METABOLISM | 0 | HALLMARK_UNFOLDED_PROTEIN_RESPONSE | 0.004 |
| HALLMARK_ADIPOGENESIS | 0 | HALLMARK_SPERMATOGENESIS | 0.005 |
| HALLMARK_REACTIVE_OXIGEN_SPECIES_PATHWAY | 0.003 | HALLMARK_GLYCOLYSIS | 0.006 |
| HALLMARK_PEROXISOME | 0 | HALLMARK_ADIPOGENESIS | 0.011 |
| HALLMARK_PI3K_AKT_MTOR_SIGNALING | 0.001 | HALLMARK_HEME_METABOLISM | 0.016 |
| HALLMARK_GLYCOLYSIS | 0.001 | HALLMARK_XENOBIOTIC_METABOLISM | 0.023 |
| HALLMARK_ALLOGRAFT_REJECTION | 0.001 | HALLMARK_PEROXISOME | 0.046 |
| HALLMARK_SPERMATOGENESIS | 0.013 | HALLMARK_ESTROGEN_RESPONSE_LATE | 0.066 |
| HALLMARK_PROTEIN_SECRETION | 0.019 | HALLMARK_ALLOGRAFT_REJECTION | 0.1 |
| HALLMARK_IL2_STAT5_SIGNALING | 0 | HALLMARK_BILE_ACID_METABOLISM | 0.099 |
| HALLMARK_MITOTIC_SPINDLE | 0.013 | HALLMARK_PANCREAS_BETA_CELLS | 0.231 |
| HALLMARK_ESTROGEN_RESPONSE_LATE | 0.008 | | |
| HALLMARK_UV_RESPONSE_UP | 0.019 | | |
| HALLMARK_ANDROGEN_RESPONSE | 0.054 | | |
| HALLMARK_CHOLESTEROL_HOMEOSTASIS | 0.08 | | |
| HALLMARK_P53_PATHWAY | 0.029 | | |
| HALLMARK_BILE_ACID_METABOLISM | 0.074 | | |
| HALLMARK_APOPTOSIS | 0.099 | | |
| HALLMARK_XENOBIOTIC_METABOLISM | 0.105 | | |
| HALLMARK_ESTROGEN_RESPONSE_EARLY | 0.197 | | |

Table S4: Gene Sets Enriched in LAMs Compared to TAMs

| Gene sets Enriched in CM TAMs vs. LAMs | | Gene sets Enriched in BC TAMs vs. LAMs | |
|---|---|---|---|
| **GS DETAILS** | **FDR q-val** | **GS DETAILS** | **FDR q-val** |
| HALLMARK_EPITHELIAL_MESENCHYMAL_TRANSITION | 0 | HALLMARK_TNFA_SIGNALING_VIA_NFKB | 0 |
| HALLMARK_ANGIOGENESIS | 0.021 | HALLMARK_TGF_BETA_SIGNALING | 0 |
| HALLMARK_COAGULATION | 0 | HALLMARK_EPITHELIAL_MESENCHYMAL_TRANSITION | 0 |
| | | HALLMARK_UV_RESPONSE_DN | 0 |
| | | HALLMARK_INFLAMMATORY_RESPONSE | 0 |
| | | HALLMARK_NOTCH_SIGNALING | 0.022 |
| | | HALLMARK_PROTEIN_SECRETION | 0.01 |
| | | HALLMARK_INTERFERON_GAMMA_RESPONSE | 0 |
| | | HALLMARK_ANGIOGENESIS | 0.058 |
| | | HALLMARK_MITOTIC_SPINDLE | 0.002 |
| | | HALLMARK_HEDGEHOG_SIGNALING | 0.067 |
| | | HALLMARK_IL6_JAK_STAT3_SIGNALING | 0.026 |
| | | HALLMARK_APICAL_JUNCTION | 0.016 |
| | | HALLMARK_KRAS_SIGNALING_UP | 0.01 |
| | | HALLMARK_COMPLEMENT | 0.025 |
| | | HALLMARK_KRAS_SIGNALING_DN | 0.031 |
| | | HALLMARK_INTERFERON_ALPHA_RESPONSE | 0.1 |
| | | HALLMARK_HYPOXIA | 0.054 |
| | | HALLMARK_WNT_BETA_CATENIN_SIGNALING | 0.197 |
| | | HALLMARK_ANDROGEN_RESPONSE | 0.149 |
| | | HALLMARK_P53_PATHWAY | 0.128 |
| | | HALLMARK_APOPTOSIS | 0.188 |
| | | HALLMARK_MYOGENESIS | 0.193 |
| | | HALLMARK_IL2_STAT5_SIGNALING | 0.205 |

Table S5: Gene Sets Enriched in TAMs Compared to LAMs
